# Detecting cell-of-origin and cancer-specific methylation features of cell-free DNA from Nanopore sequencing

**DOI:** 10.1101/2021.10.18.464684

**Authors:** Efrat Katsman, Shari Orlanski, Filippo Martignano, Ilana Fox-Fisher, Ruth Shemer, Yuval Dor, Aviad Zick, Amir Eden, Iacopo Petrini, Silvestro G. Conticello, Benjamin P. Berman

## Abstract

The Oxford Nanopore (ONT) platform provides portable and rapid genome sequencing, and its ability to natively profile DNA methylation without complex sample processing is attractive for clinical sequencing. We recently demonstrated ONT shallow whole-genome sequencing to detect copy number alterations (CNA) from the circulating tumor DNA (ctDNA) of cancer patients. Here, we show that cell-type and cancer-specific methylation changes can also be detected, as well as cancer-associated fragmentation signatures. This feasibility study suggests that ONT shallow WGS could be a powerful tool for liquid biopsy, especially real-time medical applications.

## Introduction

Circulating cell-free DNA (cfDNA) can reveal informative features of its tissue of origin including somatic genome alterations, DNA modifications, and cell-type specific fragmentation patterns (*1*). DNA methylation is a promising cfDNA biomarker and is in widespread testing as a cancer screening tool (*2*). DNA methylation can also be used to detect turnover of damaged cells in time-sensitive conditions such as myocardial infarction, sepsis, and COVID-19 (*3–6*). These studies used bisulfite-based approaches to profile methylation, but alternative approaches include immunoprecipitation (*7*) and enzymatic conversion (*8*) techniques.

Accurate calling of DNA methylation from native DNA Oxford Nanopore (ONT) sequencing has matured and now produces single base-pair resolution results highly similar to bisulfite sequencing (*9, 10*). The ONT platform is portable and has a low cost of setup, and a rapid sequencing workflow that can enable real-time medicine (*11, 12*). Native ONT sequencing requires no complex sample processing steps and no PCR amplification, making it attractive for clinical tests. Bisulfite approaches in particular involve significant degradation and loss of input material. For these reasons, whole-genome sequencing (WGS) using the ONT platform is appealing relative to other whole-genome approaches.

Due to their low cost, targeted bisulfite PCR (including multiplexed NGS versions (*13*)) are also popular for clinical methylation sequencing, and these would not require the native modification calling capability of Nanopore. However, shallow whole-genome approaches that capture multiple genomic features could potentially be more informative, especially with regard to the course of the disease (*8, 14, 15*).

ONT has primarily been used for long-read sequencing, but recent work by our group and others has shown that it can be adapted for short fragments without additional processing steps (*16–18*). As an added benefit, the ability to capture much longer cfDNA fragments than short-read sequencing may lead to new discoveries or biomarkers, as was demonstrated recently in the case of longer fragments during pregnancy (*19*).

Here, we perform a feasibility study of ONT sequencing for circulating tumor DNA (ctDNA) detection by comparing methylation and several fragmentation features to matched Illumina samples and comparable Illumina-based datasets. While our sample size is too small to determine the limits of sensitivity and specificity, we find that both cell-type specific and cancer-specific features can be reliably detected in most of our samples. The results provide confidence for pursuing this approach in larger studies.

## Results

### Estimating cell type fractions from cfNano

We first performed cell type deconvolution of healthy plasma cfDNA using DNA methylation data from either published WGBS datasets or our cfNano samples. For the external WGBS datasets, we used the methylation fractions (beta values) that were provided in the published data files. For our cfNano, we performed direct modification calling using the Megalodon software provided by ONT (https://github.com/nanoporetech/megalodon). To perform deconvolution, we used 1,000-2,000 marker CpGs per cell type based on a previously published atlas of purified cell types (“MethAtlas”, (*5, 13*)), and estimated cell type fractions using Non-Negative Least Squares (NNLS) regression as described in (*5*). In order to better understand the impact of the relatively low sequencing depth of our cfNano samples (∼0.2x genome coverage), we first performed deconvolution of all samples using downsampling experiments starting with full sequence depth down to 0.0001x genome coverage (Figure 1A and Supplementary Figures 1-3). Healthy plasma WGBS samples were taken from a recent study of 50-100x genomic coverage ((*13*), Figure 1A left “Fox-Fisher” samples), and another WGBS study with 0.5-1x coverage ((*20*) Figure 1A middle “Nguyen” samples). Finally, healthy cfNano samples were analyzed (Figure 1A right “this study”). From full depth down to 0.2x (about 2.5M aligned fragments), all samples were dominated by the expected cell types: monocytes, lymphocytes, megakaryocytes, neutrophils/granulocytes, and sometimes hepatocytes (*5*). Cell type proportions became significantly degraded at 0.05x coverage and below (corresponding to less than 700,000 aligned fragments). The common cell types were consistently found across the 23 healthy individuals in the Fox-Fisher dataset, the 3 healthy individuals in the Nguyen dataset, and the 7 healthy individuals in the cfNano dataset, both at full depth (Figure 1B) and when downsampled to 0.2x depth (Figure 1C). The same was found when cell type groups such as lymphocytes were broken down into the 25 individual types (Supplementary Figure 4A-C). Notably, a slight epithelial fraction was identified in some of the Fox-Fisher samples at 0.2x which did not appear at full 80x depth, suggesting a small but measurable amount of noise at the 0.2x coverage level.

**Figure 1:**
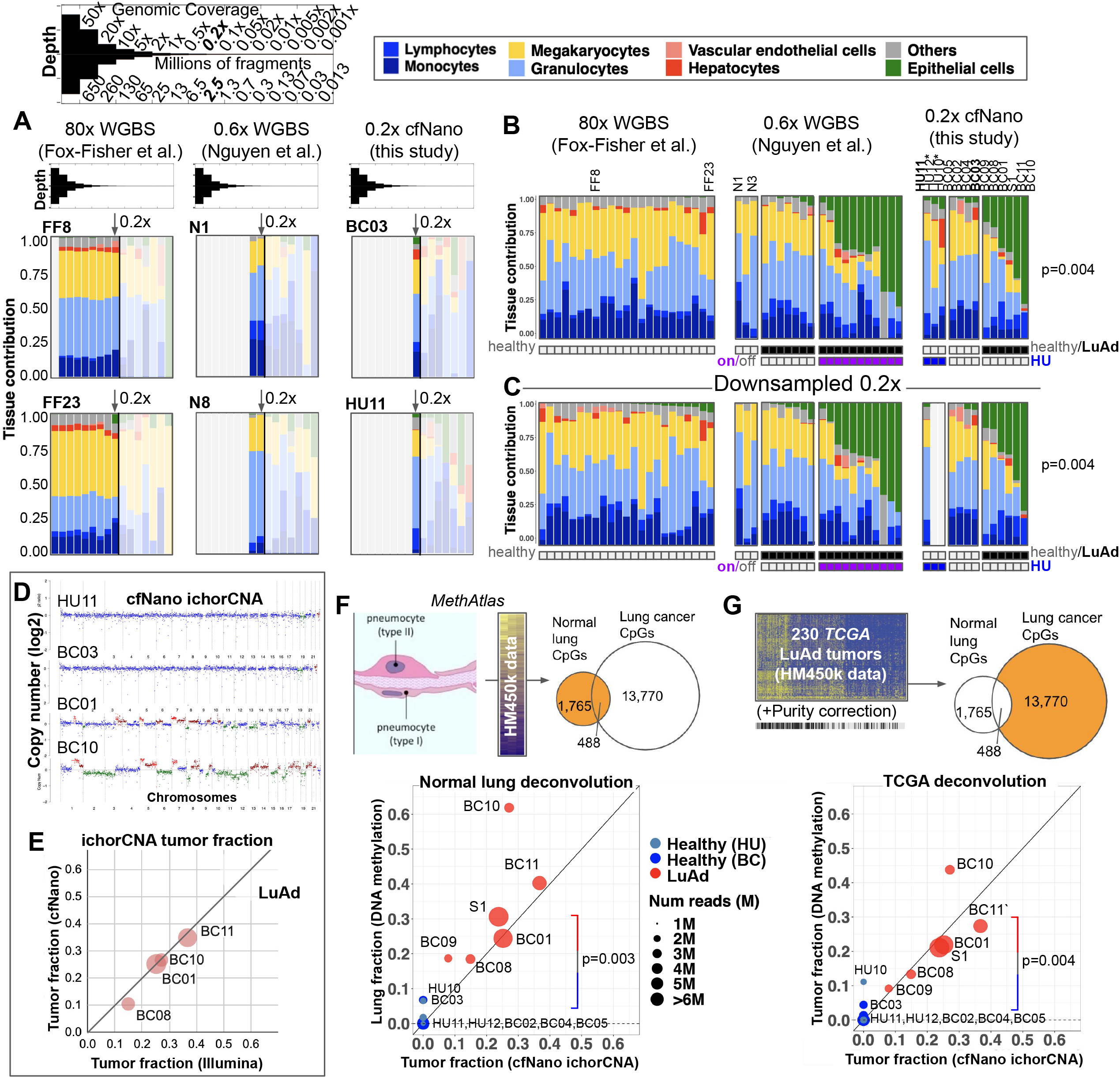
Estimating cell type fractions from cfNano. (A) Non-Negative Least Squares regression based on (*5*) was used to deconvolute cell types in healthy plasma cfDNA samples from three whole-genome DNA methylation studies. Two representative samples are shown for each study (FF8 and FF23 for the Fox-Fisher et al. study, N1 and N8 for the Nguyen et al. study, and BC03 and HU11 for our cfNano samples.) Each sample is downsampled from full read depth down to an average genome coverage of 0.001 (corresponding to approximately 13,000 fragments). All samples are shown in Supplementary Figures 1-3. (B) Deconvolution of all samples at full depth, with samples ordered within each group by epithelial cell fraction. Healthy vs. lung adenocarcinoma (LuAd) is shown as an annotation bar, as is the “on-target/off-target” status of the Nguyen et al. samples and the source site (HU Israel vs. BC Italy) for the cfNano samples. Asterisks mark the two HU samples with coverage less than 0.2x sequence depth. Statistical significance (p-value=0.004) is shown for percent epithelial in healthy cfNano samples vs. LuAd cfNano samples. (C) The same samples downsampled to 0.2x sequence depth. (D) ichorCNA CNA plots for 4 representative cfNano samples, two healthys and two LuAds. Plots for all samples are included as Additional Data File 1. (E) Tumor Fraction estimates (TF) from four LuAd samples based on ichorCNA from cfNano and matched Illumina WGS. (F) Two-component DNA methylation deconvolution of lung fraction using CpGs from MethAtlas purified lung epithelia samples, showing scatter plot of ichorCNA estimates vs. deconvolution estimates for all cfNano samples. Statistical significance is shown for DNA methylation estimate of healthy cfNano vs. LuAd cfNano samples (p-value=0.003). (G) Two-component DNA methylation deconvolution of lung cancer fraction using CpGs from TCGA LuAd tumor samples, showing scatter plot of ichorCNA estimates vs. deconvolution estimates for all cfNano samples (healthy vs. LuAd p-value=0.004). Statistical significance for panels B,C,F, and G was determined by one-tailed t-test.

The healthy cfNano individuals were divided into two groups based on source site, with one being collected and sequenced in Italy (“BC” samples) and one in Israel (“HU” samples). Despite the HU samples being lower coverage (two were between 0.10-0.15x depth), they displayed relatively similar cell type proportions (Figure 1B-C and Supplementary Figure 3).

In addition to healthy individuals, the Nguyen WGBS dataset and our cfNano dataset also contained individuals being treated for lung adenocarcinoma (LuAd), marked as “LuAd” in Figure 1B-C. In the Nguyen WGBS study, samples were collected at the time of acquired resistance to EGFR-inhibitors, and were divided into those that acquired resistance mutations in EGFR itself (labeled “on” for on-target) vs. those that acquired amplifications in alternative oncogenes MET/ERBB2 (labeled “off” for off-target). The epithelial cell fraction was much higher in the on-target patients, while the off-target patients had very low or no epithelial fraction (Figure 1B), consistent with the absence of CNAs in the off-target samples in the original study (*20*). The 6 LuAd samples in our cfNano study had similarly high epithelial fraction (Figure 1B), which was significantly higher than in the healthy patients (p=0.004). In all WGBS and cfNano samples, full depth results were highly similar to 0.2x downsampled results (Figure 1C and Supplementary Figures 1-3). Interestingly, while the Nguyen et al. study interpreted the normal-like methylation levels of the “off-target” tumors as a difference in cancer methylation patterns, our deconvolution results strongly suggest that it is due to the relative amount of cancer DNA circulating in the blood.

The fraction of cancer cells in cfDNA (“tumor fraction”) can be estimated from somatic copy number alterations (CNAs) using the ichorCNA tool (*21*), for cancers that contain a sufficient degree of aneuploidy. We estimated tumor fraction for our cfNano samples and four matched Illumina WGS samples from LuAd patients (Figure 1D, Supplementary Table 1, and Supplementary Data File 1). While the Illumina samples were sequenced at significantly higher depth (median 1.3x), the tumor fraction estimates were highly similar between cfNano and Illumina sequencing (Figure 1E). Interestingly, the ichorCNA tumor fractions were more similar to the high-depth Illumina samples than the Illumina samples themselves were when downsampled to the same depth as the cfNano samples (Supplementary Figure 5A).

To compare ichorCNA tumor fraction estimates to methylation-based estimates, we designed a “two-component” deconvolution method based on NNLS regression that used 2,253 CpGs with differential methylation between sorted lung epithelia and healthy plasma. This was based on the same array-based MethAtlas samples from (*5*) as the full deconvolution (Figure 1F). 330-1,526 of these CpGs were covered by each cfNano sample (Supplementary Table 1), which were the CpGs used for NNLS deconvolution. These DNA methylation based estimates of lung fraction and the ichorCNA estimates of cancer cell fraction were largely in agreement (Figure 1F, bottom), with all the six LuAd samples having significantly higher lung fraction compared to the seven healthy plasma samples (p=0.003). Two LuAd cases were markedly higher in the methylation-based than the CNA-based estimate (BC09 and BC10). While we have no independent data to determine which was the more accurate estimate, we hypothesize that the discrepancy may be due to either whole-genome doubling (WGD) events that are not detected by ichorCNA (WGD occurs in 297/503 or 59% of LuAd tumors from the TCGA project (*22*)), or damage to normal lung cells surrounding the tumor which die and shed their DNA into circulation (*23*).

To verify the robustness of methylation-based deconvolution, we used a mutually exclusive set of 13,770 CpGs that could distinguish TCGA LuAd tumors from healthy plasma, but were not found in the normal lung epithelia set (Figure 1H). Before applying the NNLS regression, we corrected the methylation levels of the TCGA LuAd samples to correct for their degree of non-cancer contamination (“purity correction”), since most TCGA LuAd samples contain a significant fraction of leukocytes. After this correction, the tumor fraction estimates of our cfNano samples were highly similar to those based on normal-lung specific CpGs (Figure 1H, bottom), despite the fact that the two CpG sets were completely non-overlapping. One HU healthy sample (HU005.10) had a higher lung fraction estimate than one of the cancer samples, possibly because this was the cfNano sample with the lowest sequencing coverage (0.11x). However, the methylation-based tumor fraction was still significantly higher in the LuAd samples than healthy controls (p=0.004).

We performed all deconvolution analysis using a second, and older, base modification caller (DeepSignal, (*24*)). While Megalodon called 10-20% more CpGs, the majority of CpGs called were in common between the two methods and had identical methylation states (Supplementary Figure 6A). Both the full cell-type deconvolution (Supplementary Figure 6B) and the two-component deconvolution (Supplementary Figure 6C) were highly similar between the two callers.

### Genomic context of DNA methylation changes detected using cfNano

The deconvolution analysis above was based on unannotated differentially methylated regions. In order to investigate the genomic context of lung cancer specific DNA methylation, we analyzed one hypomethylation feature associated with cell of origin (lineage-specific transcription factor binding sites) and one associated specifically with transformation (global hypomethylation). For the TFBS analysis, we identified 5,974 predicted TFBS that were specific to lung epithelia based on a single-cell ATAC-seq atlas of open chromatin within lung and other primary human tissues (*25*). In that study, adult lung tissues from multiple donors contained a strong cluster of lung pneumocyte-specific open chromatin regions (“Pal” cluster). This cluster was most strongly enriched for the binding motif for the transcription factor NKX2-1, which is a master regulatory transcription factor in this cell type (*26*). NKX2-1 activity is also known to be highly restricted to this cell type (*27*), and NKX2-1 binding sites were also the most enriched within lung adenocarcinoma ATAC-seq sites in an independent study (*28*). Because open chromatin regions are almost universally hypomethylated, we hypothesized that the 5,974 predicted NKX2-1 TFBS in lung pneumocytes would be specifically hypomethylated in healthy lung tissues and in lung tumors. We confirmed this using WGBS data from TCGA (*29*) (Supplementary Figure 7A).

We next plotted plasma cfDNA methylation levels at these same predicted NKX2-1 sites from the published Illumina WGBS studies and our cfNano study (Figure 2A). In healthy samples from three WGBS studies and our own cfNano samples, NKX2-1 sites were fully methylated. In contrast, the LuAd samples from both the Nguyen et al. WGBS study (Figure 2A, middle) and our cfNano study (Figure 2A, right) had substantial demethylation. In both studies demethylation could only be observed in the higher tumor fraction samples (“on-target” samples in the WGBS study, and samples with ichorCNA TF>0.15 in the cfNano study). As a negative control, we selected predicted TFBS from a cell type not expected to be found either in healthy plasma or LuAd. We used the adrenal cortical cluster (“Adc” cluster) from (*25*), which was highly enriched for the KLF5 binding motif. These sites were fully methylated in plasma samples from both healthy and LuAd individuals (Supplementary Figure 7B). cfNano profiles were nearly identical using DeepSignal methylation calling (Supplementary Figure 7C).

**Figure 2:**
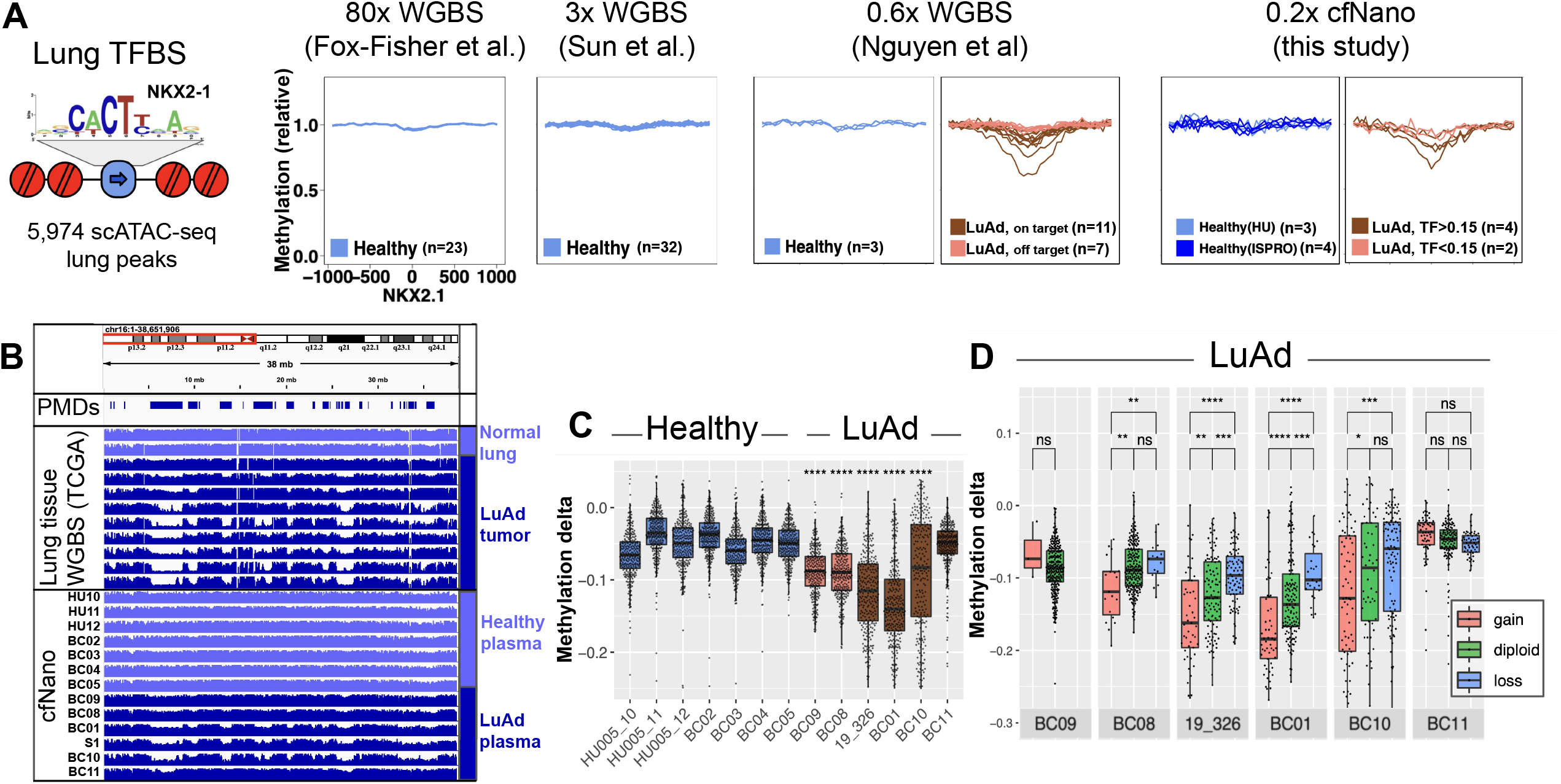
Genomic context of DNA methylation changes detected using cfNano. (A) Plasma cfDNA methylation levels were averaged from -1kb to +1kb at 5,974 pneumocyte-specific NKX2-1 transcription factor binding sites (TFBS) taken from (*25*). All methylation values are fold change relative to the flanking region (region from 0.8kb-1kb from the TFBS). From left to right, plots show 23 healthy plasma samples from (*13*), and 32 healthy plasma samples from (*42*), 3 healthy and 18 LuAd WGBS samples from (*20*), and 7 healthy and 6 LuAd cfNano samples from this study. (B) Average DNA methylation across chr16p, comparing lung tissue WGBS (top) to plasma cfNano samples from this study (bottom). Reference Partially Methylated Domains (PMDs) are taken from (*29*). (C) Methylation delta is shown for all 10Mbp bins overlapping a reference PMD (methylation delta defined as the average methylation of the bin minus the average methylation genome-wide). Each cancer sample was compared to the group of healthy samples using a one-tailed t-test, and statistical significance is shown using asterisks. (D) 10Mbp PMD bins were stratified by copy number status for each cancer sample using ichorCNA, and statistically significant differences were calculated by performing one-tailed Wilcoxon tests within each sample.*p<0.05, **p<0.01, ***p<0.001, ****p<0.0001.

Global DNA hypomethylation is one of the hallmarks of the cancer epigenome. It has long been proposed as a general marker for circulating tumor DNA (*30*), and this was recently verified for lung cancer using shallow plasma cfDNA WGBS (*20*). We have shown that this “global” hypomethylation is not completely global, and occurs preferentially within large domains called Partially Methylated Domains (PMDs) and specifically at CpGs with a local sequence context termed “solo-WCGW” (*29*). We replotted WGBS methylation data from TCGA normal lung and lung tumor tissues, showing a typical chromosome arm where strong hypomethylation occurs within the PMD regions identified in (*29*) in the cancer samples (Figure 2B, top). In our cfNano samples, strong hypomethylation was also found exclusively in the cancer samples in the same PMD regions (Figure 2B, bottom). To quantify this genome-wide, we plotted the methylation change (relative to the sample-specific whole-genome average) of PMD solo-WCGW CpGs within each 10 Mbp genomic bin that overlapped a common PMD region from (*29*) (Figure 2C). As expected, five of the six cancer samples were significantly hypomethylated relative to the healthy controls (p<0.0001). Overall, there was significantly more hypomethylation across all cancer sample bins (mean=-0.10,SD=0.07) than across all healthy sample bins (mean=-0.05,SD=0.04), corresponding to a p-value<2.2e-16 by one-sided Wilcoxon test. In the final LuAd case (BC11), no PMD hypomethylation could be detected (Figure 2C). This is not surprising given the high degree of variability associated with global hypomethylation in cancer (*29*), a process that is not entirely understood but is affected both passively through mitotic divisions as well as actively by dysregulation of several chromatin modifiers (*31, 32*).

Reasoning that copy number altered regions would have skewed proportions of tumor-derived DNA and thus different levels of PMD hypomethylation, we divided the PMD bins based on the copy number status of each sample from ichorCNA. In four of the five cases with significant hypomethylation overall, the amplified bins had significantly more hypomethylation than diploid regions (Figure 2D). In the one remaining case (BC09), there were not enough PMDs with CNAs for an accurate measurement. Conversely, deleted showed significantly less hypomethylation than diploid regions, although this trend only reached statistical significance in two cases. In the future, the combined analysis of CNAs and global hypomethylation may provide a stronger cancer-specific signal than each feature alone. PMD hypomethylation profiles were nearly identical using DeepSignal methylation calling (Supplementary Figure 7D-F).

### cfNano preserves nucleosome positioning signal

Cell-free DNA circulates primarily as mononucleosomal fragments, and mapping the positions of these mononucleosomes can be used to identify cell-type specific positioning (reviewed in (*1*)). CTCF binding sites provide a good test of whether these signals are detectable, since they eject a central nucleosome and position 10 phased nucleosomes on either side of their binding site (*33*) (Figure 3A). Around a set of 9,780 CTCF binding sites, cfNano mononucleosome locations recapitulated this expected pattern (Figure 3B, top), which was identical to the pattern based on matched Illumina WGS of greater sequence depth (Figure 3B, bottom). These were also identical when both cfNano and Illumina libraries were downsampled to an equal number of 2M fragments (Figure 3C). The CTCF binding site also demethylates CpGs located approximately 200bp of on either side (*33*), and this DNA methylation pattern was also recapitulated in our cfNano samples (Supplementary Figure 8).

**Figure 3:**
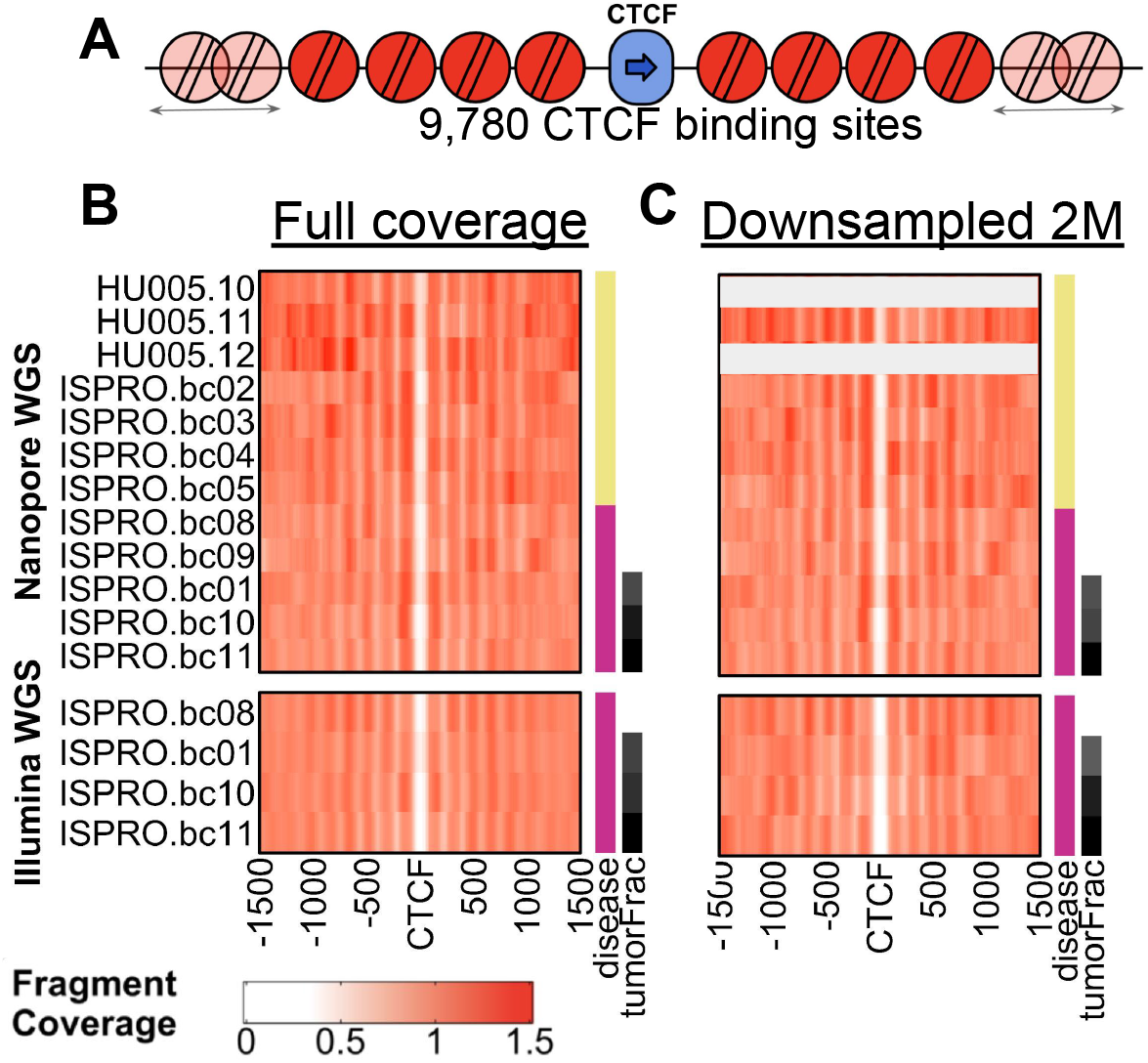
cfNano preserves nucleosome positioning signal. (A) Alignments to 9,780 CTCF motifs within non-promoter ChIP-seq peaks were taken from (*33*). (B) Sequence coverage of mononucleosomes (130-155bp) from cfNano samples is shown as fold-change vs. average coverage across the genome (top). Mononucleosome coverage for matched Illumina samples (bottom). (C) Same analysis, using a randomly selected downsampling of 2 million reads from each sample. Two cfNano samples with less than 2M reads total are omitted.

We tried the same mononucleosome mapping approach for the 5,974 lung-specific NKX2-1 TFBS from Figure 2A. While the demethylation signal was detectable (Figure 2A), we could not detect any mononucleosome positioning signal (data not shown). Lung-specific nucleosome positions would only be present on a fraction of the fragments, so the signal from these fragments may be masked by those from non-lung cell types. But given that the inherent nucleosome positioning information is present (as shown by the CTCF example), more advanced normalization and quantification techniques may reveal these cell-type specific fragments more sensitively in the future.

### Cancer-associated fragmentation length features of cfNano vs. Illumina WGS

Specific fragment lengths have been associated with cancer-derived cfDNA fragments (reviewed in (*1*)) and these have been used as accurate cancer classifiers (*15*). Specifically, shorter mononucleosome fragments (<150bp) tend to be enriched for cancer-derived fragments (*34*). Density plots of fragment length showed that our cfNano cancer samples were enriched in these short mononucleosome fragments relative to healthy controls (Figure 4A). We used the definition from (*34*) and (*15*) to calculate the ratio of short mononucleosomes (100-150bp) to all mononucleosomes (100-220bp). The short mononucleosome ratio was significantly higher in the high tumor fraction cancer cases (mean=0.24, SD=0.03) than in the healthy cases (mean=0.16, SD=0.03), which corresponded to a t-test p-value of p=0.038 (Figure 4B). We compared these ratios calculated from our cfNano libraries with those calculated from the matched Illumina WGS libraries which were available for four of the six cancer samples, and the two library types were strongly correlated (Figure 4C). We hypothesized that improvements to Nanopore basecalling could improve alignment and adapter trimming, so we also compared base calling done with the Guppy basecaller at the time of sequencing (“2019” version) to the new “high accuracy calling” basecalling (“HAC”) performed on all samples in 2022. The new ratios with the new basecalling were slightly more similar to the matched Illumina libraries (Figure 4C).

**Figure 4:**
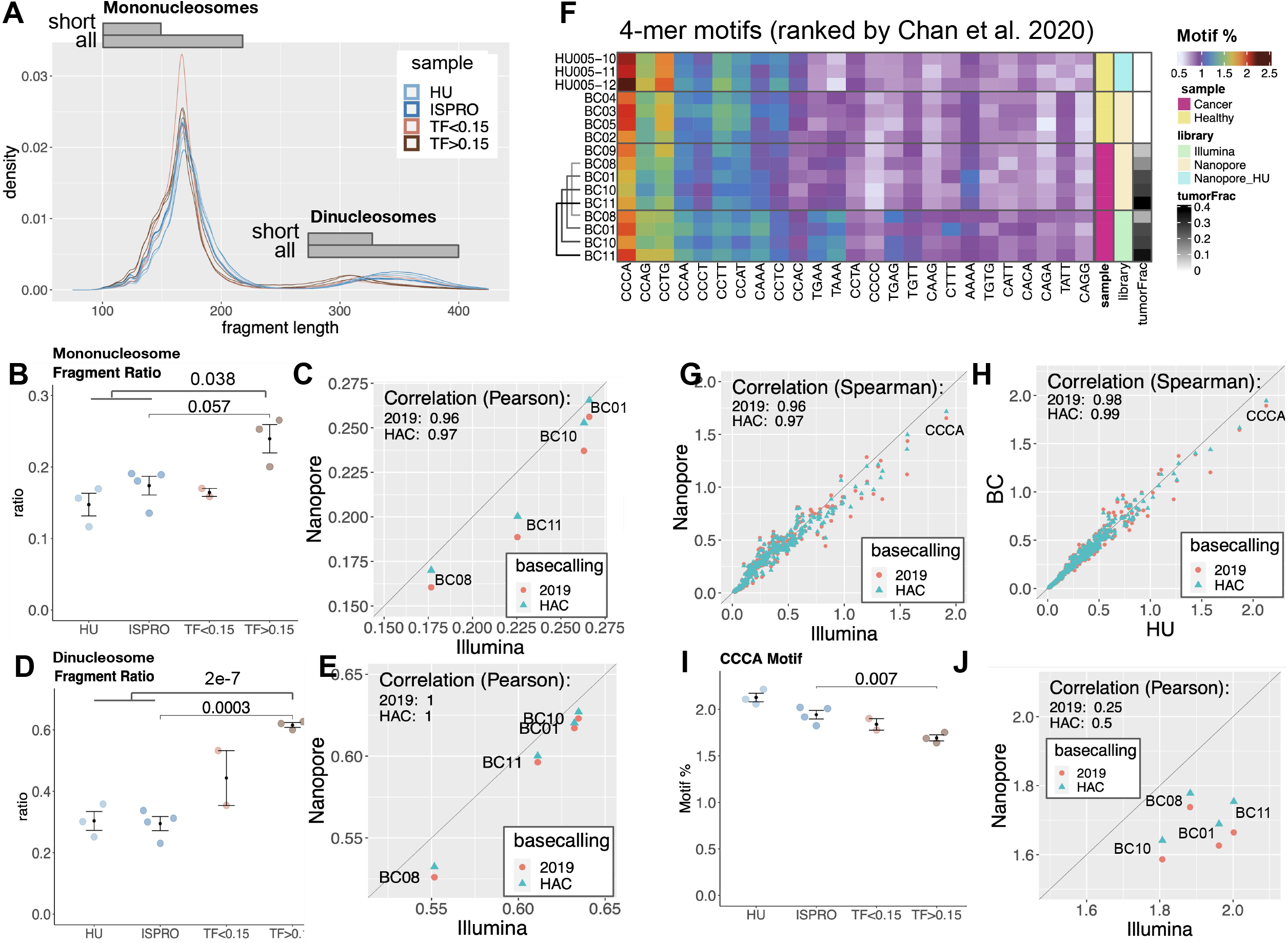
Cancer-associated fragmentation features of cfNano vs. Illumina WGS. (A) Fragment length density plot for each cfNano sample, with cancer samples divided into low tumor fraction (TF<0.15) and high tumor fraction (TF>0.15) based on ichorCNA. Short mononucleosomes are defined as 100-150bp (*15, 34*) and short dinucleosomes are defined at 275-325bp. (B) The ratio (fraction) of short mononucleosome fragments (100-150bp) to all mononucleosome fragments (100-220bp). (C) Short mononucleosome ratios based on cfNano are compared to short mononucleosome ratios based on matched Illumina WGS libraries for four LuAd cases. cfNano samples were processed with either 2019 Oxford Nanopore Fast basecalling model (2019) or 2022 Oxford Nanopore High Accuracy model (HAC), as indicated by color. (D) The ratio (fraction) of short dinucleosome fragments (275-325bp) to all dinucleosome fragments (275-400bp). (E) Short dinucleosome ratios based on cfNano vs. Illumina WGS ratios for matched LuAd samples. (F) Frequency of 4-mer sequences occurring at fragment ends, for cfNano vs. matched Illumina samples. The 25 most frequent 4-mers are shown in ranked order based on frequencies in (*36*). (G) End motif frequencies for all 256 possible 4-mers, comparing average frequency in four cfNano samples vs four matched Illumina WGS samples. (H) End motif frequencies, comparing average frequency in four healthy HU Italy cfNano samples vs three healthy HU Israel cfNano samples. (I) Frequency of CCCA 4-mer in all cfNano samples. (J) CCCA 4-mer frequencies from cfNano samples vs. frequencies calculated from Illumina WGS for four matched LuAd samples. Statistical significance levels for panels B,D, and I were determined by two-tailed t-test.

While they have not been studied as extensively as mononucleosomes, ref. (*34*) also showed that dinucleosomes were significantly shorter in cancer fragments than non-cancer fragments. This is clear from the density plots of our cfNano samples (Figure 4A), so we used the size range suggested by ref. (*34*) to calculate the ratio between short dinucleosomes (275-325bp) and all dinucleosomes (275-400bp). The short dinucleosome ratio showed even more separation between cancer vs. healthy cfNano samples than the mononucleosome ratio, with the high tumor fraction cancer cases (mean=0.62, SD=0.01) than in the healthy cases (mean=0.30, SD=0.04), corresponding to a t-test p-value of p=2E-7 (Figure 4D). We compared dinucleosome ratios calculated from our cfNano libraries with those calculated from the matched Illumina WGS libraries, and they were nearly perfectly correlated (Figure 4E). When we looked across all samples, the short mononucleosome ratio was highly correlated with the short dinucleosome ratio (Supplementary Figure 9D). Interestingly, this correlation held across the healthy samples as well as the cancer samples, indicating that the same underlying mechanism affects circulating cfDNA from both cancer and non-cancer cell types.

One of our cfNano samples used a different (non-barcoded) adapter design method from all other libraries, and this sample was a clear outlier in fragment length (Supplementary Figure 9A-D). This reinforces the caution that should be taken when comparing fragmentomic features across different library designs. We also investigated the effect on sequencing depth on cancer-associated features, by comparing full-depth datasets with datasets created by randomly choosing 2M fragments for each library (Supplementary Figure 9E-H). Sequence depth had almost no effect on either cfNano or Illumina samples down to 2M fragments.

### Cancer-associated fragment end features of cfNano vs. Illumina WGS

The four bases immediately flanking cfDNA fragmentation sites have biased sequence composition that differs between cancer-derived and non-cancer-derived fragments (*35, 36*). To study this in our cfNano samples, we first plotted the 25 most abundant 4-mer end motifs that were identified in an Illumina-based study of healthy plasma cfDNA (*36*), using a heatmap to indicate motif frequencies in each of our cfNano and matched Illumina samples (Figure 4F). There was broad agreement across all samples, although some differences between cfNano and Illumina libraries were clearly noticeable. When we plotted average Nanopore vs. Illumina frequencies for all 256 possible 4-mers, it appeared that the less abundant motifs had slightly higher frequencies in Nanopore, while the more abundant motifs had slightly lower frequencies in Nanopore (Figure 4G). Nevertheless, the relative frequencies were highly concordant overall (PCC=0.97). These were slightly more concordant when we used the 2022 “high accuracy” (HAC) basecalling compared to the original 2019 basecalling (PCC=0.97 vs. PCC=0.96). The degree of difference between the two batches of cfNano healthy samples (“BC” sequenced in 2019 and “HU” sequenced in 2022) showed only slight differences using the HAC basecalling (PCC=0.99).

Of particular interest is the CCCA end motif, which is typically the most abundant 4-mer in healthy plasma and its reduction was shown to be a cancer marker in several cancer types including lung cancer (*35, 36*). CCCA indeed has the highest frequency across all our cfNano and Illumina WGS samples (Figure 4F-H), and was significantly lower in our three high tumor fraction cancer samples than the healthy samples (Figure 4I). However, there was a clear difference between the healthy samples generated in the “HU” and “ISPRO” batches, which we presume to be technical since these two batches behaved similarly with respect to fragment length and methylation features. We therefore only did a direct statistical comparison within the ISPRO batch, and indeed CCCA frequency in high TF tumors (mean=1.6,SD=0.06) was significantly lower than in ISPRO healthy samples (mean=1.9,SD=0.13), leading to a t-test p-value=0.007 (Figure 4I).

We found additional 16 motifs that were as significant as CCCA, although none survived FDR adjustment and so will have to be validated in larger studies (Supplementary Table 3). Like fragment lengths, end motif frequencies were not sensitive to downsampling to 2M fragments (Supplementary Figure 10A-D). Additionally, the relative frequencies of four cfNano cancer samples were not concordant with their matched Illumina WGS libraries (Figure 4J). We conclude from this that end motifs are particularly sensitive to changes in library strategy and sequencing platform, and caution must be taken when comparing across multiple batches. This is not surprising, given that fragment representation can be skewed by a number of variables during library construction and amplification, as well as sequencing errors and downstream bioinformatic steps such as adapter trimming (in our cfNano processing, we also exclude fragment ends that are soft-clipped). End motifs are highly susceptible to these biases, because even a single base pair difference results in a completely different motif. Recent benchmarking has highlighted how error frequencies can differ by sequence context between the Nanopore and Illumina platforms (*37*).

## Discussion

While the sample size is small, our results suggest that cancer-specific features of DNA methylation, fragmentation, and CNA are broadly concordant between cfNano and Illumina-based WGS and WGBS methods. Downsampling analysis showed that the genomic coverage we targeted with cfNano (minimum of 2.5M aligned reads or 0.2x genome coverage) was sufficient to detect cancer-derived DNA in all samples based on DNA methylation. Cancer-associated fragmentomic features were not detected in all samples, but this is likely due to biological variability rather than sequence depth, based on similar studies using Illumina-based approaches (*34*), as well as our own results running the fragmentomic analysis on downsampled libraries. Notably, our results suggest that short dinucleosomes could be a more robust cancer marker than short mononucleosomes, although this will need to be validated in larger studies.

While most features agreed between cfNano and Illumina-based datasets, we identified fragment end motifs as one that was especially sensitive to sequencing platform differences. With the small sample size here, it is not possible to determine which of the two platforms provides the truer results. It is tempting to hypothesize that Nanopore provides a more accurate representation of actual fragment frequencies, since it does not include any PCR amplification. On the other hand, Nanopore error rates are higher and may lead to less accurate adapter trimming or a different spectrum of sequencing errors within the end motifs (although we did try to control for this by using the reference genome sequence rather than the read sequence, and by filtering out reads with soft-clipped ends). Despite these differences, we were able to detect the best known cancer-specific end motif (CCCA) when we constrained the analysis to samples from our main cfNano batch (“ISPRO”). We also detected slight differences in end motif frequencies between our own Illumina WGS and earlier published WGS data, and between our two cfNano batches which were sequenced two years apart. This indicates that end motif frequencies are susceptible to batch effects in general and should be analyzed cautiously between batches. In the future, proper controls should be developed to compare this feature across datasets sequenced at different times.

With the small number of metastatic cancer samples used in our cfNano study, it was not possible to define the lower limit of methylation sensitivity. However, the fact that some epithelial/lung content was found when downsampling higher coverage WGBS samples and in our cfNano healthy samples that had the lowest coverage, suggests that specificity needs to be improved at this very low coverage. We believe this could be improved significantly if whole-genome DNA methylation atlases (WGBS or similar) can be generated for individual cell types purified from human tissues. The purified cell type atlas we used here was based on the Illumina HM450k platform, which covers only about 10-15% of cell type specific methylation markers. Alternatively, cancer markers could come directly from discovery WGBS studies on cancer plasma cfDNA. Such datasets do exist in the private domain, but remain proprietary (e.g. (*2*)).

Even if deconvolution can be improved bioinformatically, we will almost certainly need higher read coverage for applications that require more sensitive detection. The cfNano samples analyzed here were all from metastatic disease and had relatively high tumor fraction (>=10%), and this does not represent the situation for cancer screening or detection of minimal residual disease. Thus, the Nanopore platform will need to be able to consistently produce tens of millions of reads to enable those applications. Using cfNano directly to discover new cancer markers would not necessarily require greater coverage, but would require large numbers of patient samples.

Several other current limitations of Nanopore should be considered. The cost per base of Nanopore sequencing is currently several-fold higher than Illumina, although the new generation of ONT PromethION sequencers are meant to address this as well as increase throughput. Single-nucleotide and indel error rates are higher for the ONT platform, which could pose an issue for whole-genome analysis of mutational signatures (*38, 39*), something we do not investigate here but is theoretically possible from cfNano. While Nanopore error rates have improved significantly over the past several years, this is a weak point that should be considered if mutations are a priority.

There are several areas where Nanopore methylation sequencing may provide unique strengths over other methods. Bisulfite-based sequencing leads to DNA fragmentation and degradation, and can obscure fragmentation patterns (*40*). Additionally, bisulfite sequencing can not distinguish between 5mC and other modifications such as 5hmC, 5fC, and 5CaC, whereas these are all in principle detectable by Nanopore. Longer cfDNA fragments have not been extensively studied due to limitations of short-read technology, and these could be valuable both for biomarker discovery and the basic biology of cfDNA (i.e. (*19*)).

Despite the limitations discussed above, the simplicity of native ONT sequencing and the number of features that can be extracted from a single run, combined with the low cost and portability of sequencer, make it an interesting proposition for clinical settings. Fast sample prep and sequencing times can allow a complete methylation analysis from sample preparation to computational classification in as little as 1-3 hours, enabling real-time medical applications in cancer (*11, 12*). Because DNA methylation can differentiate non-cancer cell types as well, Nanopore liquid biopsy could be used to monitor collateral damage to adjacent tissue in cancer (*23*), or urgent conditions in other areas of medicine such as myocardial infarction, sepsis, and COVID-19 (*4–6, 41*).

## Code availability

Source code for fragmentomic analysis is available at https://github.com/Puputnik/Fragmentomics_GenomBiol. Source code for combined CNV and PMD methylation analysis is available at https://github.com/Puputnik/CNV_Methylation_Genome_Biol_2022. Source code for methylation deconvolution is available at https://github.com/methylgrammarlab/cfdna-ont. An archive of all source code is deposited in Zenodo DOI:10.5281/zenodo.6445784.

## Data availability

Processed data files for the analyses described here are available at GEO accession GSE185307 and at Zenodo DOI: 10.5281/zenodo.6448476. These include “anonymized BAM files” generated by Megalodon for all cfNano samples. Anonymized BAM files contain all fragmentomic information as well as read-specific DNA methylation calls. We also include methylation BED files for both Megalodon and DeepSignal. Raw data files (fastq) for cfNano samples are deposited in EGAD00001006888.

## Acknowledgments

We thank Joshua Moss and Yaping Liu for critical review of the manuscript, Sara Isaac and Myriam Maoz for assistance with sample processing, and Tiago Silva and Irene Unterman for helpful discussions. We thank the Dennis Lo lab for sharing healthy control cfDNA data from EGAD00001001602, and the Le Son Tran lab for sharing healthy and LuAd plasma cfDNA data from 10.6084/m9.figshare.16817941.v1. Computation was carried out on the Hebrew University Research Computing Services cluster, and we acknowledge Yaron Weitz and Ori Adam for their help and support. Ben Berman, Efrat Katsman, and Shari Orlanski received startup support from the Hebrew University, the Kamea B program of the Israel Ministry of Aliyah and Immigrant Integration, and the Israel Cancer Research Fund Project Grant (845755). Filippo Martignano was supported by the Italian Ministry of Health (SG-2019-12370279).

## Author Contributions

BPB, FM, and SGC conceived the project. BPB, EK, SO, FM, and SGC designed the computational analyses. BPB, EK, SO, and FM performed computational and statistical analyses and generated figures. IP provided cancer plasma samples and associated clinical information. IFF, RS, and YD provided healthy plasma samples and associated clinical information. FM, AZ, and AE performed ONT sequencing. BPB wrote the manuscript, with contributions from SGC, FM, EK, and SO. BPB and SGC co-supervised the project. The first three authors (EK,SO,FM) are co-lead authors with an equal contribution to the work, and they have the right to list their names first in their CVs. The last two authors (SGC,BPB) are co-senior authors who jointly supervised the work, and they have the right to list their names last in their CV.

## Figure legends

**Supplementary Figure 1: DNA methylation deconvolution for Fox-Fisher et al. samples**. Each sample from (*13*) was downsampled from full depth to 0.001x coverage, and sample ordering is the same as Fig 1B-C. Short names are used, and full sample information is available in Supplementary Table 2.

**Supplementary Figure 2: DNA methylation deconvolution for Nguyen et al. samples**. Each sample from (*20*) was downsampled from full depth to 0.001x coverage, and sample ordering is the same as Fig 1B-C. Short names are used, and full sample information is available in Supplementary Table 2.

**Supplementary Figure 3: DNA methylation deconvolution for cfNano samples**. Each cfNano sample from the current study was downsampled from full depth to 0.001x coverage, and sample ordering is the same as Fig 1B-C. Short names are used, and full sample information is available in Supplementary Table 1.

**Supplementary Figure 4: Full cell type assignments in deconvolution analysis**. (A) Cell-type deconvolution for WGBS and cfNano datasets, using 25 cell types from (*5*). (B) 25 cell type deconvolution of all samples downsampled to 0.2x sequence coverage. (C) The four cell-type groups from Figure 1 (Lymphocyte, Granulocyte, Epithelial, and Other) and which of the 25 cell types were collapsed into each group. All cell types not assigned to one of the four groups are shown as a singleton cell type in Figure 1.

**Supplementary Figure 5: ichorCNA tumor fractions of downsampled Illumina samples**. Four Illumina plasma samples from LuAd patients are shown. ichorCNA tumor fraction was computed at full sequence depth (x axis) and by randomly downsampling the Illumina samples to have the same number of fragments as the corresponding cfNano sample.

**Supplementary Figure 6: Calling cfNano methylation with two different methods**. (A) DeepSignal and Megalodon were used to call CpG methylation for each cfNano sample. CpGs were divided into those covered by DeepSignal only, Megalodon only, or Both. Those covered by both were divided into those that got identical methylation status vs. different methylation status. (B) Grouped cell type deconvolution is shown for all samples for Megalodon and DeepSignal processed data. Megalodon version is reproduced from Figure 1B. (C) Two-component deconvolution is shown for all samples for Megalodon and DeepSignal processed data. Megalodon versions are reproduced from Figure 1F and 1G, respectively.

**Supplementary Figure 7: Genomic context of DNA methylation changes**. (A) Methylation in 18 TCGA WGBS non-lung tumors (left) and 11 TCGA WGBS lung tumors and adjacent normal tissue (right) from (*29*). Plasma cfDNA methylation levels were averaged from -1kb to +1kb relative to 5,974 pneumocyte-specific NKX2-1 transcription factor binding sites (TFBS) taken from (*25*). All methylation values are shown as relative to the flanking region (from 0.8kb-1kb relative to TFBS). (B) 9,274 adrenal cortical cell specific KLF5 TFBS taken from (*25*). From left to right, plots show 23 healthy plasma samples from (*13*) and 32 healthy plasma samples from (*42*), followed by 3 healthy and 18 LuAd WGBS samples from (*20*) and 7 healthy and 6 LuAd cfNano samples from this study (C) cfNano methylation levels for lung NKX2-1 (same as Figure 2A), using DeepSignal methylation calling. (D) cfNano methylation levels for adrenal cortical KLF5 (same as this Figure, panel B), using DeepSignal methylation calling. (E) IGV analysis (same as Figure 2B) using DeepSignal methylation calling. (F-G) Genome-wide PMD bin analysis (same as Figure 2C-D) using DeepSignal methylation calling.

**Supplementary Figure 8: cfNano preserves fragmentomic and DNA methylation markers of nucleosome positioning**. Alignments to CTCF motifs within 9,780 distal ChIP-seq peaks from (*33*). (A, top) cfDNA fragment coverage shown as fold-change vs. average coverage depth across the genome. The plot includes only fragments of length 130-155bp to maximize resolution. (A, bottom) Matched Illumina samples of higher sequencing depth (median 17.0M fragments in Illumina vs. 6.4M in ONT samples). (B) CTCF DNA methylation of Nanopore samples from this study at CTCF sites. (C) DNA methylation from seven lung tissue WGBS samples from TCGA (*29*).

**Supplementary Figure 9: Effects of downsampling on fragment length of cfNano and Illumina WGS**. (A-C) Data from Figure 4A,B,D are reproduced with the addition of sample 19_326 (which used a different, non-barcoded, cfNano adapter design), as well as matched Illumina samples. (D) Short mononucleosome ratios (x axis) plotted against short dinucleotide ratios (y axis). Panels (E-H) show the same plots as panels A-D, but with each sample randomly downsampled to 2M fragments. Statistical significance levels for panels B,C,F, and G were determined by two-tailed t-test.

**Supplementary Figure 10: Effects of downsampling on fragment end features of cfNano and Illumina WGS**. (A-B) are reproduced from main Figure 4F and 4I, with the addition of sample 19_326 (which used a different, non-barcoded, cfNano adapter design), as well as matched Illumina samples. Panels (C-D) show the same plots, but with each sample randomly downsampled to 2M fragments.Statistical significance levels for panels B and D were determined by two-tailed t-test.

**Supplementary Table 1:** cfNano sample information and statistics

**Supplementary Table 2:** External WGBS sample information

**Supplementary Table 3:** Tumor vs. normal differences for 4-mer end motifs

**Supplementary Data File 1:** ichorCNA plots for all cfNano and matched Illumina WGS samples.

## Methods

### ISPRO Plasma cfDNA samples, library construction, and sequencing

ISPRO Samples, library construction and sequencing were described in our initial publication of these sequences (*17*). The original sample names from that study are listed in Supplementary Table 1. Notably, one sample (S1/19_326) was produced using a different library kit (SQK-LSK109 vs. NBD-EXP104+SQK-LSK109 for all other samples). This is the singleplex library kit, which results in shorter adapter-ligated templates overall (due to the lack of barcodes) and thus responds differently to the equivalent clean up bead concentration. Also adapter trimming is performed differently in 19_326 due to the library kit differences. For these reasons, fragmentomic properties are not directly comparable between 19_326 and other samples. We thus analyzed 19_326 separately for all fragmentomic analyses (included in Supplementary Figures 9-10), but included it in all figures when analyzing methylation and copy number alterations, where small differences in fragment length are not expected to make a difference. Standard MinKNOW runtime control was used without modification (S1 using distribution version 18, and all others using version 19).

### HU Plasma cfDNA samples, library construction, and sequencing

HU healthy samples are cfDNA extracted from 4mL plasma as described in (*13*). The original sample names from that study are listed in Supplementary Table 1. Barcoded libraries were created using the NBD-EXP104 and SQK-LSK109 kits as for ISPRO samples. They were sequenced on a single flow cell, using standard MinKNOW runtime control (distribution v.21.11.7) without modification.

### 2019 Fast model basecalling and alignment of Nanpore fast5 files

Basecalling was done using MinKNOW distribution v.19.06.9 Guppy version 3.0.6+9999d81. For singleplex sample (S1/19_326), adapters were trimmed with Porechop (https://github.com/rrwick/Porechop) using: “--discard_middle --extra_end_trim 0”. For both HU and ISPRO samples, alignments were performed to GCF_000001405.39_GRCh38.p13 with minimap2 (Version 2.13-r850), using the parameters “-ax map-ont --MD”.

### 2022 High Accuracy Calling (HAC) basecalling and alignment of Nanopore fast5 files

HU and ISPRO Fast5 files were demultiplexed with guppy_barcoder (Version 5.0.16+b9fcd7b5b) using “--trim_barcodes --barcode_kits EXP-NBD104”. Individual barcode Fast5 files were processed into fastq and bam files using Megalodon v. 2.4.2 with the following command-line parameters “--edge-buffer 0 --mod-min-prob 0 --guppy-params ‘-d /usr/local/hurcs/guppy/6.0.1/data -- barcode_kits EXP-NBD104 --trim_barcodes’ --remora-modified-bases dna_r9.4.1_e8 hac 0.0.0 5mc CG 0 --guppy-config dna_r9.4.1_450bps_hac.cfg”. Internally, Megalodon used Guppy server version 6.0.1+652ffd1, and basecalling model r9.4.1_450bps_hac. By default, Megalodon filters out multi-mapping (supplementary) reads and uses the minimap2 “map-ont” mode to filter low quality mappings.

Individual tile Fast5 files were run individually, and mod_mapping.bam files were merged using samtools merge (v1.14). Samtools/HTSlib versions before v.1.14 do not handle the Mm/Ml modification tages. Fastq files were merged by concatenating (cat) all individual basecalls.fastq files. Merged mod_mapping.bam files have been deposited in GEO GSE185307 and at Zenodo DOI: 10.5281/zenodo.6448476. Because Megalodon reports only the *reference* sequence in the BAM records, and does not report any base substitutions, these are anonymous BAM files which do not contain any SNP information, and thus contain no personally identifiable information. These are the primary files used for both fragmentomic and methylation analysis, described in more detail below, and are available from GEO GSE185307 and at Zenodo DOI: 10.5281/zenodo.6448476.

### DeepSignal methylation calling and processing

We used DeepSignal Version 0.1.8 (4), with model “model.CpG.R9.4_1D.human_hx1.bn17.sn360.v0.1.7+/bn_17.sn_360.epoch_9.ckpt”, which was downloaded from the DeepSignal Google Drive (https://drive.google.com/open?id=1zkK8Q1gyfviWWnXUBMcIwEDw3SocJg7P). We used the DeepSignal call_mods (modification_call) output tsv file, extracting the (strand-specific) methylation calls for each CpG from column 9 (called_label field), and calculated a methylation beta value by taking the number of methylated reads (value 1) divided by the total number of reads (value 0 or value 1). These were collapsed into a bedgraph file with a value between 0-1 for every CpG covered. These are available as file “grouped-beta-value_bedgraph.zip” in GEO accession GSE185307 and at Zenodo DOI: 10.5281/zenodo.6448476. All genomic coordinates are in GRCh38 and are zero-based.

### Megalodon methylation calling and processing

Basic running of Megalodon is described above. To extract (stranded) methylation information from the mod_mapping.bam files, we used modbam2bed (https://github.com/epi2me-labs/modbam2bed) v.0.4.5, specifying a minimum probability threshold of 0.667, and filtering out positions with 0 confident reads using awk. The full command line was “modbam2bed --cpg -t 4 -a 0.333 -b 0.667 | awk ‘($5>0){print} > out.bed”. All coordinates are in GRCh38 and are 0-based. These files are named “*.5mC.cut0.667.hg38.bed.gz”. Column 11 corresponds to the percent of reads methylated. Modbam2bed does not provide a column for the actual number of reads that this percentage is based on, but it can be calculated from the other columns. `readCount=(col5*col10)/1000`. We also profide a simple bedgrpah with just the methylation fraction (beta) values in files named “*cut0.667.hg38.sorted.bedgraph.gz”. These can be loaded into any genome browser. Both file types are available in GEO accession GSE185307 and at Zenodo DOI: 10.5281/zenodo.6448476

### Mapping of cfNano methylation data to HM450k probes

Using the zero-based stranded bed files from modbam2bed (“5mC.cut0.667.hg38.bed.gz” files), we mapped each CpG covering either the forward or reverse strand of each CpG on the Infinium 450k array. For each modbam2bed stranded column, we first got the readCount as `(col5*col10)/1000`. We then multiplied the methylation percentage by the read count for each strand to get the number of methylated reads. Then we divided the sum of the methylated read counts by the sum of the total read counts to get the unstranded percent methylation (beta value).

### Methylation calling from external WGBS datasets

For Fisher-Fox et al., methylation “beta.gz” files were obtained from GSE186888, and processed as recommended using wgbs_tools (https://github.com/nloyfer/wgbs_tools) beta2bed function to obtain fraction methylated and read count for each CpG. For Nguyen et al., bed files with methylation fractions and read counts were obtained from Figshare 10.6084/m9.figshare.16817941.v1.

For Sun et al., we obtained fastq files from EGAD00001001602 and aligned using Biscuit (https://github.com/huishenlab/biscuit) v.0.3.15.20200318 using the command line “biscuit align -t 16 hg38.fa CTR153.fq.gz -b 1” piped into samblaster (*43*) v.0.1.24 to mark and remove duplicates with the command line “samblaster -i stdin -o stdout -M --excludeDups -- addMateTags --ignoreUnmated -d CTR153.hg38_discordant.sam -s CTR153.hg38_split.sam -- maxSplitCount 2 --maxUnmappedBases 50 --minIndelSize 50 --minNonOverlap 20 –u CTR153.hg38_.fastq --minClipSize 20”.

### Methylation coverage downsampling

To downsample methylation coverage from bed files with read count and fraction methylated columns, we used a custom Perl script in the https://github.com/methylgrammarlab/cfdna-ont repository called downsampleMethylBed.pl. This script treats each read at each CpG as an independent observation, and then randomly samples from these until it has enough observations to reach the average genomic coverage requested. To obtain the coverage levels shown in Figure 1, it was run with the command line “downsampleMethylBed.pl --coverageLevels 1E-3,2E-3,5E-3,1E-2,2E-2,5E-2,1E-1,2E-1,5E-1,1E0,2E0,5E0,1E1,2E1,5E1,8E1 --fracTotalFieldsFrom0 -3,4 --ncpgsGenome 28217005”.

### Full cell type methylation deconvolution

For the full cell type deconvolution in Figures 1A-C, we used the non-negative least squares regression (NNLS) method from (*5*). Specifically, we used the code from https://github.com/nloyfer/meth_atlas/blob/master/deconvolve.py. For Megalodon methylation analysis, we used the top 1,000 hypermethylated and 1,000 hypomethylated CpGs for each of the 25 cell types provided. Full results were plotted using a modified version of the original deconvolve.py which we have deposited here: https://github.com/methylgrammarlab/cfdna-ont/blob/main/deconvolution_code/deconvolution_moss/plot_deconv.py. These are shown in Supplementary Figure 4A-B.

For Figure 1A-C, we collapsed cell types into 8 groups using the file https://github.com/methylgrammarlab/cfdna-ont/blob/main/deconvolution_code/deconvolution_moss/group_file_for_plot_green_epithilial.csv (shown visually in Supplementary Figure 4C). We plotted results using code in https://github.com/methylgrammarlab/cfdna-ont/blob/main/deconvolution_code/deconvolution_moss/deconvolution_plot.R. For DeepSignal methylation data, the procedure was the same except we used the top 2,000 hypermethylated and top 2,000 hypomethylated CpGs, to account for the significantly smaller number of CpGs called in the DeepSignal data (shown in Supplemental Figure 6A).

### ichorCNA analysis

Samtools (Version 1.9) was used to filter BAM alignments, unmapped reads, secondary and supplementary reads, reads with mapping quality less than 20 as in (*44*), and reads longer than 700bp. For Illumina alignments we trimmed all ‘N’ nucleotides from the 3’ ends of fastq data, alignments were performed to GCF_000001405.39_GRCh38.p13 with BWA mem (43), duplicates were marked using picard MarkDuplicates and removed with samtools;read pairs without the properly-paired flag were removed.Pipelines used for preprocessing and filtering of both Nanopore and Illumina data are available at https://github.com/Puputnik/Fragmentomics_GenomBiol. Somatic copy number analysis was performed using the ichorCNA package v.0.3.2 (*21*). We used default settings to determine copy number alterations and tumor fraction for each cancer sample. If the percentage of genome covered by CN alterations was less than 15%, then the tumor fraction was determined to be unstable and set to 0. ichorCNA tumor fraction estimates are available in Supplementary Table 1, and genomic CNA plots for all samples are available as Additional Data File 1.

### Two-component cell type methylation deconvolution using healthy lung epithelia

To determine lung fraction specifically from different datasets, we devised a “two-component” version of the NNLS regression model described above. To compose the atlas of differentially-methylated probes in 25 human tissues and cell types, we used the data collected and tissue-specific feature selection method from the MethAtlas package (https://github.com/nloyfer/meth_atlas)(*5*). The script feature_selection.m was used to select Lung_cell epithelial specific CpGs. For the Megalodon version, the cutoff was set to select the top 1,000 hypermethylated and the top 1,000 hypomethylated probes, for the three Lung_cell epithelia samples vs. the four healthy plasma cfDNA samples from (*5*). For DeepSignal methylation data, the procedure was the same except we used the top 2,000 hypermethylated and top 2,000 hypomethylated CpGs, to account for the significantly smaller number of CpGs called in the DeepSignal data (shown in Supplemental Figure 6A). We removed any probe that did not have valid (non-NA) values for 2 or more of the Lung_cell samples and 2 or more of the healthy plasma samples.

For each probe, the 450k beta values were averaged to produce a single Lung-specific beta value *<u>X</u>*<u>_l_</u>. The same was done for the four plasma cfDNA samples from to yield a healthy cfDNA beta value *<u>X</u>*<u>_2_</u>. We used the Lawson-Hanson algorithm for non-negative least squares (NNLS) (https://cran.r-project.org/web/packages/nnls) to perform non-negative least squares regression as in (*5*). Specifically, we identified non-negative coefficients *β*_1_and *β*_2_, representing the fraction of Lung cells and normal blood cells in the Nanopore cfDNA mixture, respectively, subject to the constraints *argmin*_*β*_||*Xβ* ≥ *Y*||_2_ and *β* ≥ 0. Then a single Lung fraction *β* was determined by having *β*_1_ and *β*_2_ sum to 1, with the equation 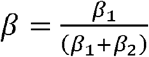.

### Two-component cell type methylation deconvolution using TCGA lung tumors

We downloaded the Infinium 450k beta value files for TCGA Lung Adenocarcinoma (LUAD) tumors using the ELMER packaged in Bioconductor (*45*). We removed any probe that did not have valid (non-NA) values for 2 or more of the LUAD samples and 2 or more of the healthy plasma samples. In order to make this analysis completely independent from the healthy lung epithelia deconvolution analysis, we excluded 488 that were in the 2,000 probe set for the Megalodon healthy lung analysis, and an additional 396 that were in the 4,000 probe set for the DeepSignal healthy lung analysis (described above). We then performed a t-test to compare the methylation beta values of these LUAD specific probes to the four plasma cfDNA samples from the MethAtlas paper (*5*), requiring a Benjamini-Hochberg corrected FDR of <0.001 and an absolute beta value difference of 0.3 or greater.

### Correcting TCGA methylation model for cancer cell purity

NNLS was performed as above for the TCGA lung tumor deconvolution, with the following adaptation. The deconvolution assumes that each of the reference cell types is a representation of the purified cell type, but this is not the case for bulk TCGA tumors which have a median of luekocyte fraction of 30% (*22*). For each probe in each TCGA cancer sample, we corrected for this by solving for the equation *M*_*m*_ = *M*_*c*_*β* + *M*_*l*_ (1 − *β*), where *M*_*m*_ is the methylation of the mixture, *M*_*c*_ is (unknown) methylation of the cancer cells, *M*_*l*_ is the (known) methylation of the leukocytes, and *β* is the (known) percentage of cancer cells in the mixture. *M*_*l*_ was taken as the average of white blood cell samples from the MethAtlas (*5*), and *β* was taken as the “tumor purity” estimate based on somatic copy number alterations from the TCGA PanCan Atlas project using the ABSOLUTE program (*22*), downloaded from the PanCan Atlas website (TCGA_mastercalls.abs_tables_JSedit.fixed.txt, URL https://gdc.cancer.gov/about-data/publications/pancanatlas). We used the pure cancer cell estimates M, and performed NNLS regression as described above.

### DNA methylation in 10 Mbp PMD bins

To generate Figures 2D-E, GRCh19 segmentation results from our previous CNV analysis (*17*) were divided into non-overlapping 10Mb bins. Copy number status of each bin was determined by log2ratio segment mean > 0.10 and < -0.10 for Gain and Loss respectively. For the healthy samples, 10Mb bins were generated from the whole genome. GRCh38 Methylation files were converted to GRCh37 using liftover R package. We selected only the bins overlapping one or more common Partially Methylated Domains (PMDs) from (*29*). Within these PMD bins, we took the average of all “solo-WCGW” CpGs overlapping a PMDs, with “solo-WCGW” annotation also from (*29*). We calculated the bin average from these CpGs as sum(frac_methylation_each_CpG)/CpG_count. We then subtracted this bin average and subtracted it from the average of all CpGs in the genome, to get the Methylation Delta shown in Figure 2C-D. Common PMD and solo-WCGW annotations were taken from file https://zwdzwd.s3.amazonaws.com/pmd/solo_WCGW_inCommonPMDs_hg38.bed.gz. Statistical significance between each cancer sample’s bins and all pooled healthy sample bins in Figure 2D was calculated by one-sided Wilcoxon test (because we decided a priori to look only for hypomethylation in the cancer samples). For copy number analysis in Figure 2D, each pair of copy number groups was compared using a one-sided Wilcoxon test, to test the hypotheses that diploid should have higher methylation than amplified regions, and deleted regions should have the highest methylation. The files and pipeline used for this analysis (including segmentation results from Martignano et al. (1)) are available at https://github.com/Puputnik/CNV_Methylation_Genome_Biol_2022.

### Transcription factor binding site (TFBS) analysis

First, we used HOMER to identify predicted NKX2-1 binding sites (using the HOMER built in matrix “nkx2.1.motif”) across the GRCh38 genome, and removed any site within the ENCODE blacklist. For normal lung cell analysis, we intersected this list with 6,754 ATAC-seq peaks identified in the pneumocyte (PAL) cluster 13 CREs from (*46*) (downloaded from supplemental table 6 of that paper “Table_S6_Union_set_of_cCREs.xlsx”). We then selected 5,974 peaks that overlapped a predicted NKX2-1 TFBS, and centered each on the predicted NXK2-1 TFBS. If multiple TFBS were present in the peak, we took the motif with the highest HOMER log-odds match score. This TFBS set is available as file “nkx2.1.incluster13_distalPeaks_PAL.bed.highestScoreMotifs.hg38.bed” in GEO accession GSE185307 and at Zenodo DOI: 10.5281/zenodo.6448476. To calculate relative methylation levels, raw methylation levels in each bin were divided by the mean methylation within all bins from -1000 to -800 and +800 to +1000 across all NKX2-1 sites. For Supplementary Figure 7A, we used all WGBS cancer types that were represented by normal tissues in the scATAC-seq atlas, as this was the atlas used to define pneumocyte specific (PAL) peaks. For TCGA lung and non-lung samples in Supplementary Figure 7A, we downloaded TCGA WGBS bedgraph files from https://zwdzwd.github.io/pmd (*29*). We used all WGBS cancer types that were represented by normal tissues in the scATAC-seq atlas, as this was the atlas used to define pneumocyte specific (PAL) peaks. These TGCA types included LUAD and LUSC (Lung tissue from atlas), CRC (Transverse colon tissue from atlas), BRCA (Breast tissue from atlas), STAD (Stomach tissue from atlas), and UCEC (Uterus tissue from atlas).

### *KLF5 transcription factor binding site (TFBS) analysis* (Supplementary Figure 7B)

As with *NKX*.*2* above, we used HOMER to identify predicted KLF5 binding sites (using the HOMER built in matrix “klf5.motif”) across the GRCh38 genome, and removed any site within the ENCODE blacklist. As a control we intersected this list with 9,274 ATAC-seq peaks identified in the cluster 43 CREs from (*46*) (downloaded from supplemental table 6 of that paper “Table_S6_Union_set_of_cCREs.xlsx”). We then selected 1,762 peaks that overlapped a predicted KLF5 TFBS, and centered each on the predicted KLF5 TFBS. If multiple TFBS were present in the peak, we took the motif with the highest HOMER log-odds match score. This TFBS set is available as file “klf5.incluster43Distal.txt.highestScoreMotifs.bed” in GEO accession GSE185307.

### CTCF nucleosome positioning analysis

We used 9,780 evolutionarily conserved CTCF motifs occurring in distal ChIP-seq peaks, which were taken from (*33*). Nanopore or Illumina fragments within the size range of 130-155bp were used for fragment coverage analysis, with reads being extracted from BAMs as described above. These shorter mononucleosomal fragments showed similar nucleosomal patterns but gave higher spatial resolution than 156-180 bp fragments. Deeptools (Version 3.5.0) bamCoverage was used with the parameters “--ignoreDuplicates -- binSize -bl ENCODE_blacklist -of bedgraph --effectiveGenomeSize 2913022398 -- normalizeUsing RPGC”. For Illumina WGS, we used the additional parameter “--extendedReads 145”. The bedgraph was converted to a bigwig file using bigWigToBedGraph downloaded from UCSC Genome Browser. This bigwig file was passed to Deeptools computeMatrix with the command line parameters “reference-point --referencePoint center -out table.out”, and the table.out file was imported into R to create fragment coverage heatmap.

### Fragment length analysis

Samtools (Version 1.9) was used to filter BAM alignments, unmapped reads, secondary and supplementary reads, reads with mapping quality less than 20 as in (*44*), and reads longer than 700bp. For Illumina alignments we trimmed all ‘N’ nucleotides from the 3’ ends of fastq data, alignments were performed to GCF_000001405.39_GRCh38.p13 with BWA mem (43),duplicates were marked using picard MarkDuplicates and removed with samtools.Pipelines used for preprocessing and filtering of both Nanopore and Illumina data, and analyzed data are available at https://github.com/Puputnik/Fragmentomics_GenomBiol. In addition, only reads with barcodes at both ends (obtained using the -- require_barcodes_both_ends flag while demultiplexing) were used for fragment length analysis of the multiplexed samples (all except 19_326). Reads with soft clipping at either the 5’ or 3’ ends were removed. Fragment length was calculated from the Minimap2 BAM CIGAR column by summing all counts. Short mononucleosome ratio was calculated as 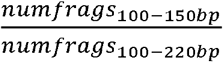 (150bp is the same cutoff for short fragments used in (*34*)). Short dinucleosome ratio was calculated as 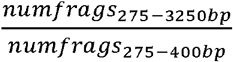 (this was determined visually from Figure 2D of publication (*34*)).

### End motif analysis

Fragments and reads were processed and filtered as in fragment length analysis. We only used end 1 of both cfNano and Illumina reads, to avoid problems with adapter trimming at end 2. To avoid biases that would affect end motif analysis, we also removed reads with any soft clipping at end 1. The first 4 bases of each fragment were extracted and used for 4-mer analysis. To avoid errors in Nanopore base calling, these 4 bases were extracted from the reference genome. Motif frequency was calculated as 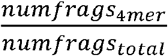. For top 25 motifs and ranking order in Figures 4 and Supplementary Figure 10, we used (*36*). Files and pipelines used for Fragment length and end motif analyses are available at https://github.com/Puputnik/Fragmentomics_GenomBiol

### Statistical tests

Student’s t-test for all sample comparisons where at least one test group had less than five samples, otherwise Wilcoxon test was used.

