## Supplementary figures and images for "Detecting cell-of-origin and cancer-specific methylation features of cell-free DNA from Nanopore sequencing"

### ichorCNA-Illumina-downsamp-cfNano.pdf

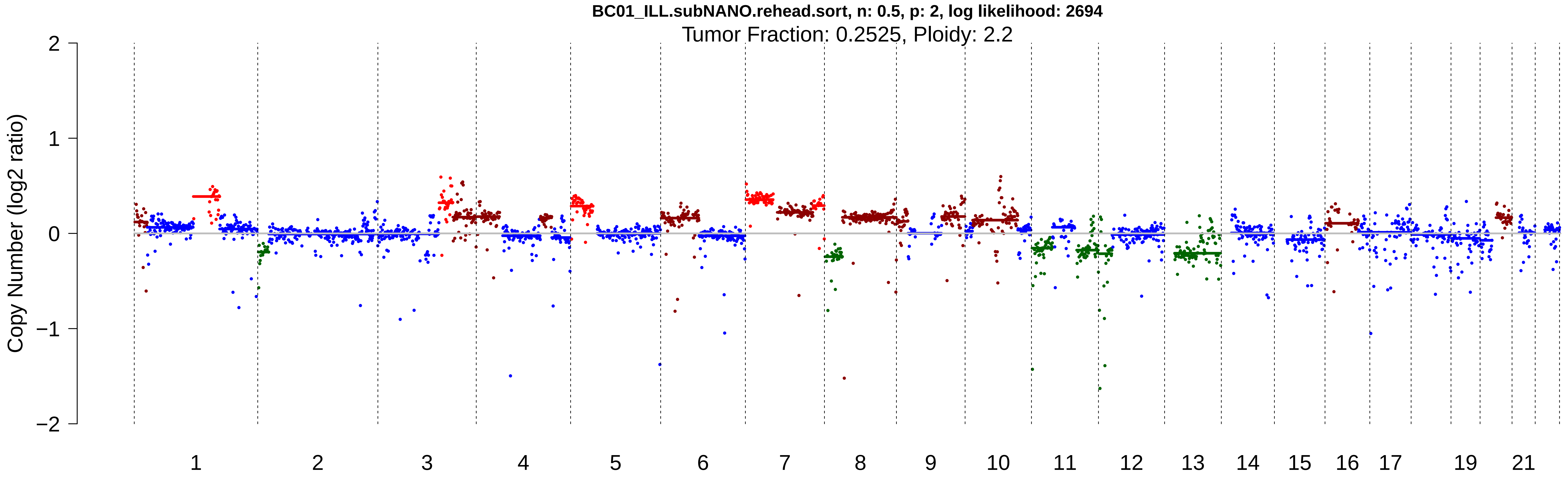

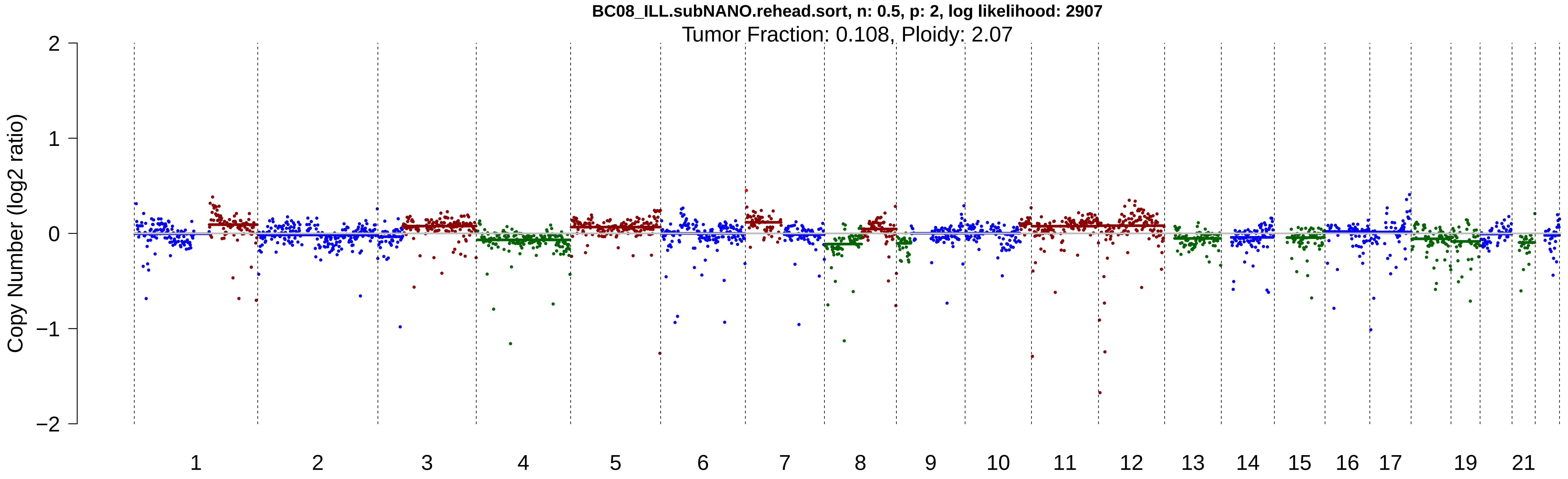

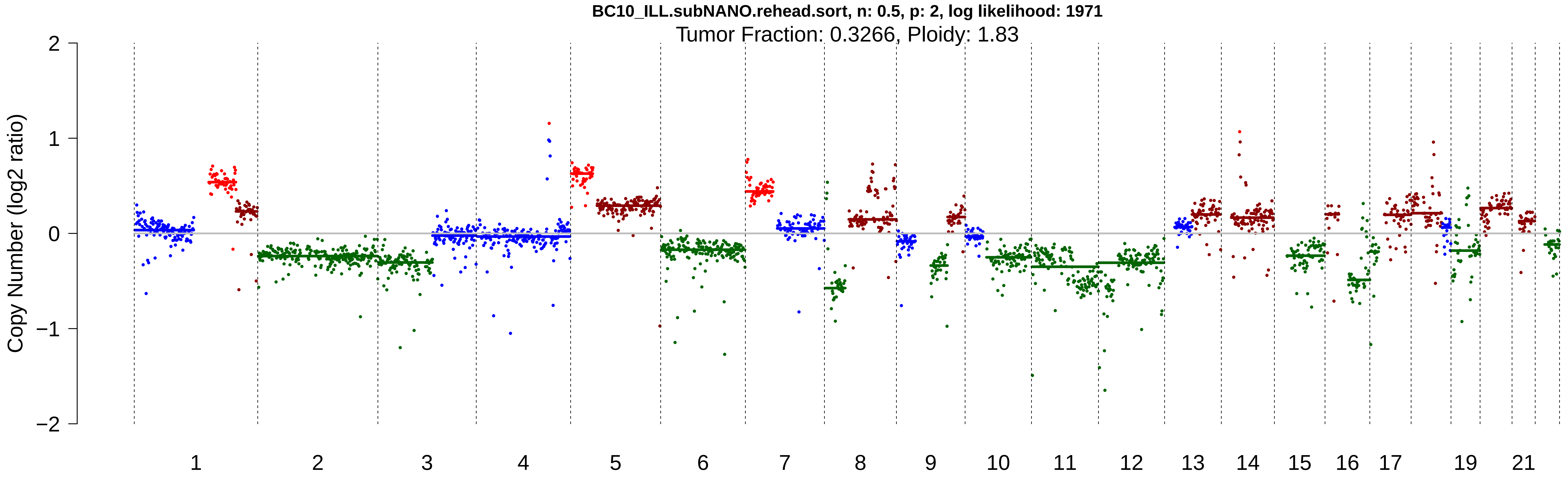

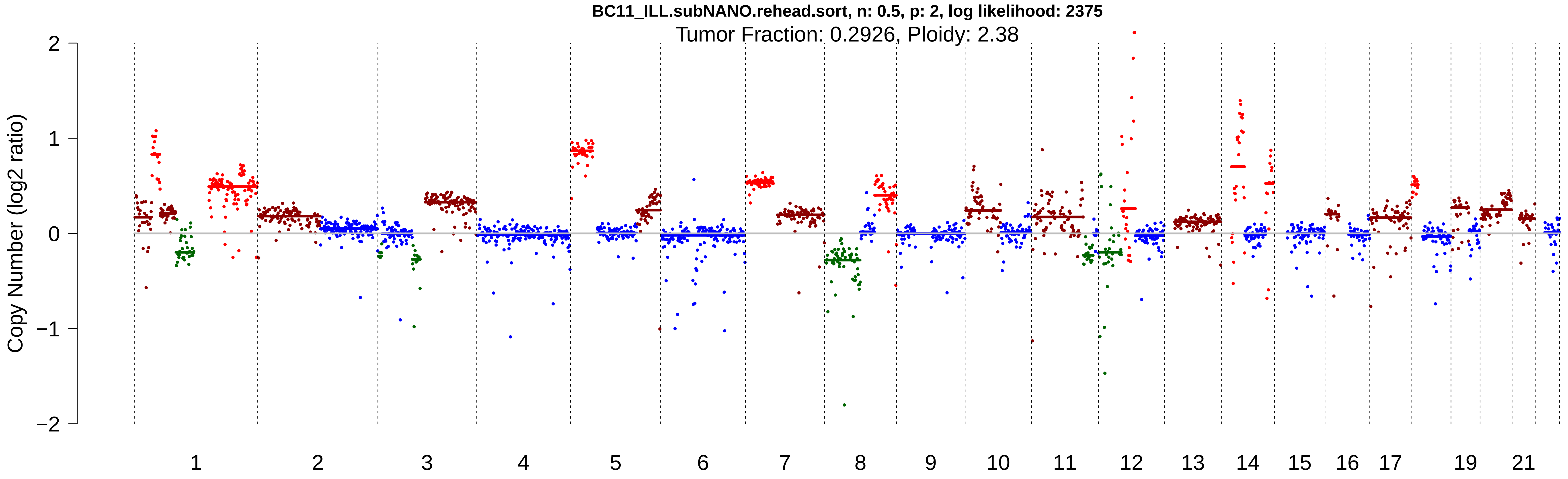

### ichorCNA-Illumina-original.pdf

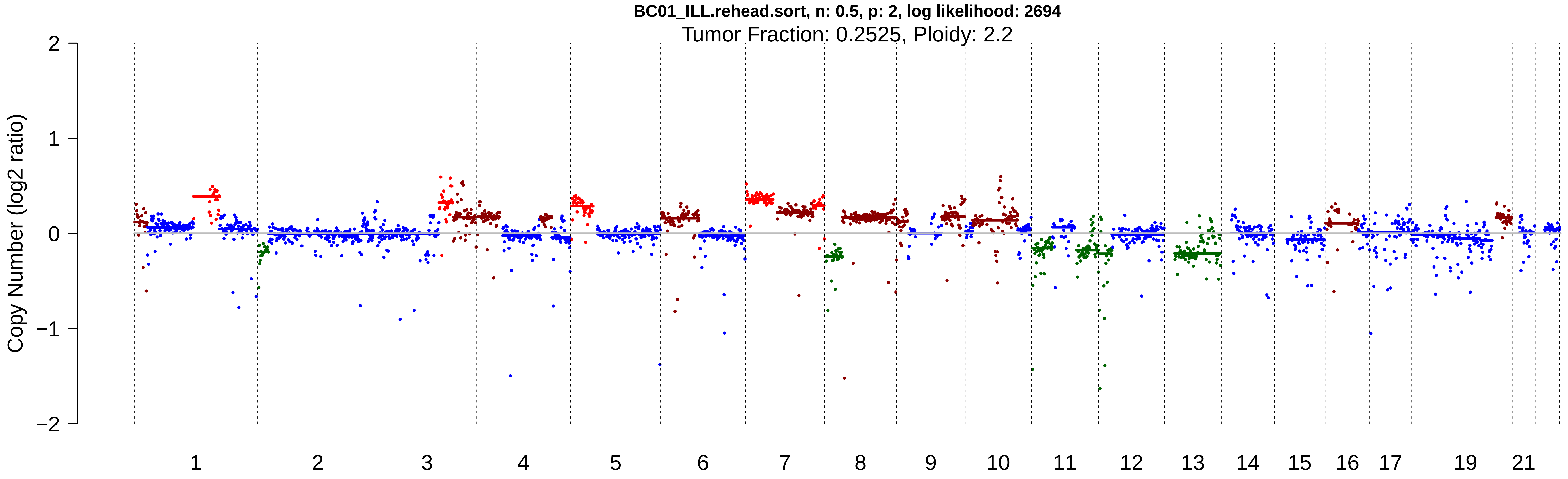

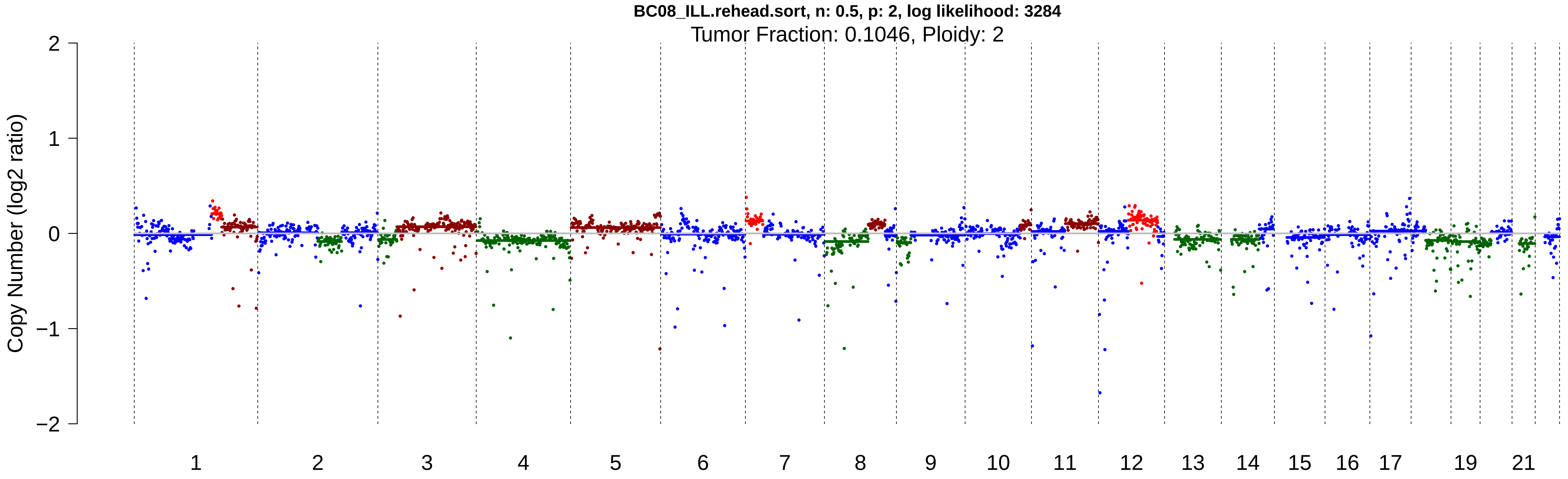

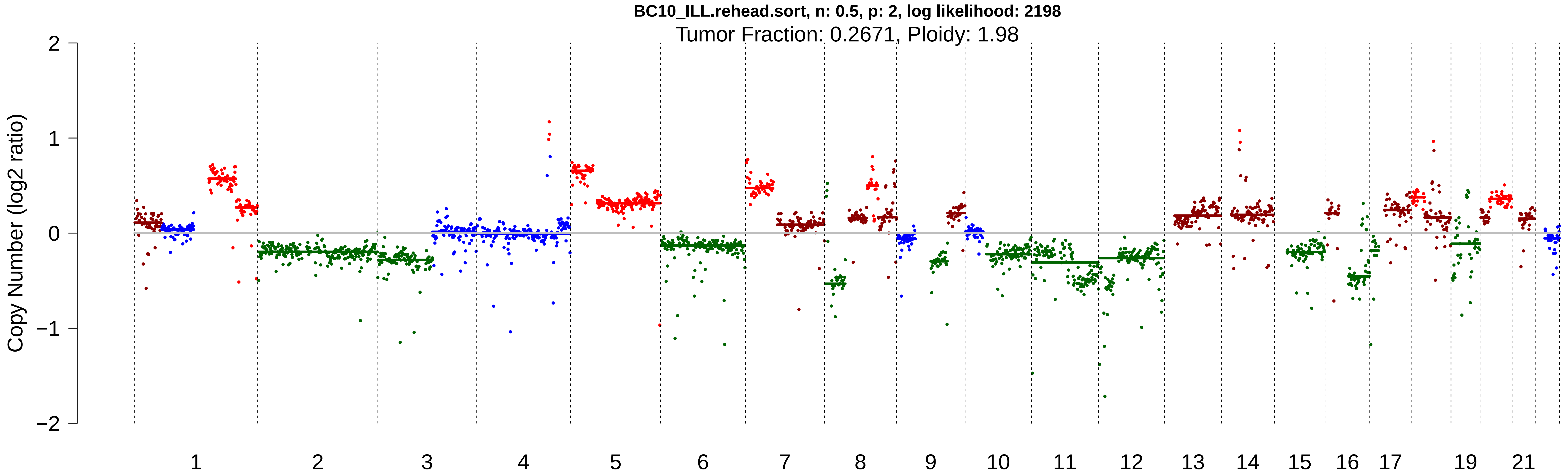

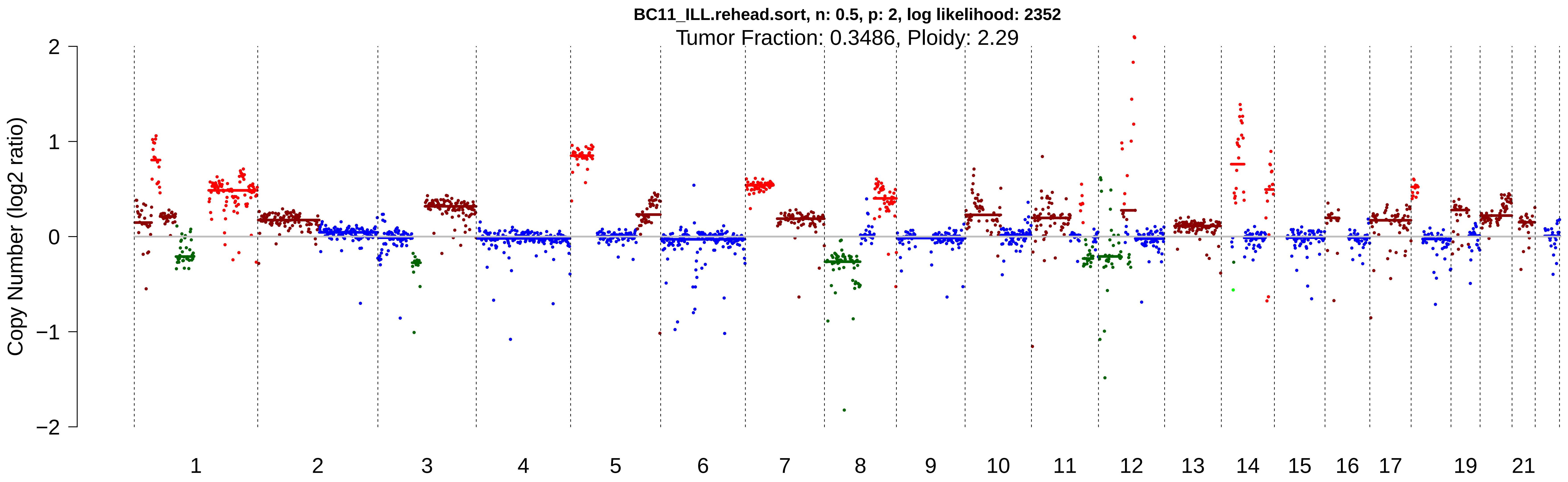

### Supplementary Figures 1-10

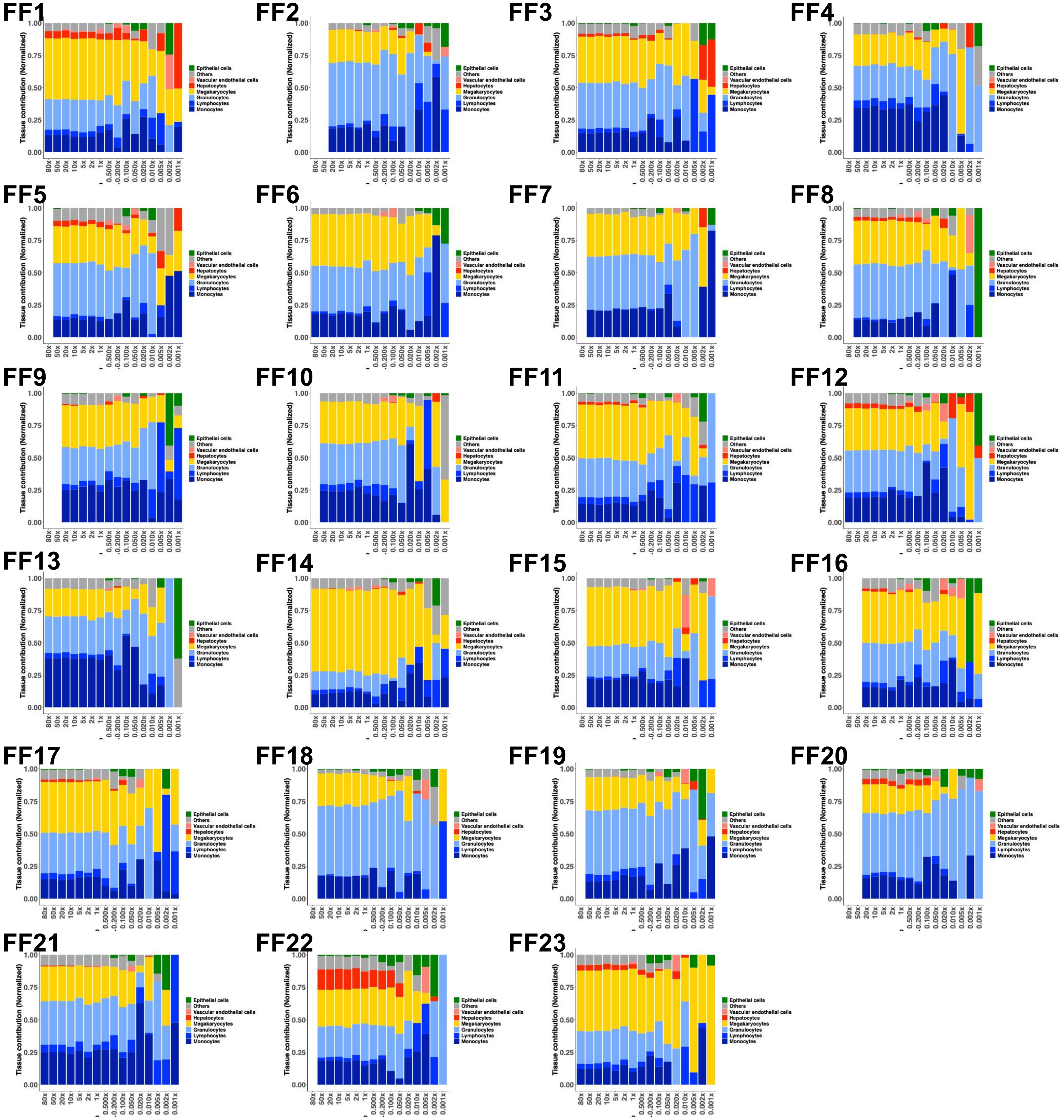

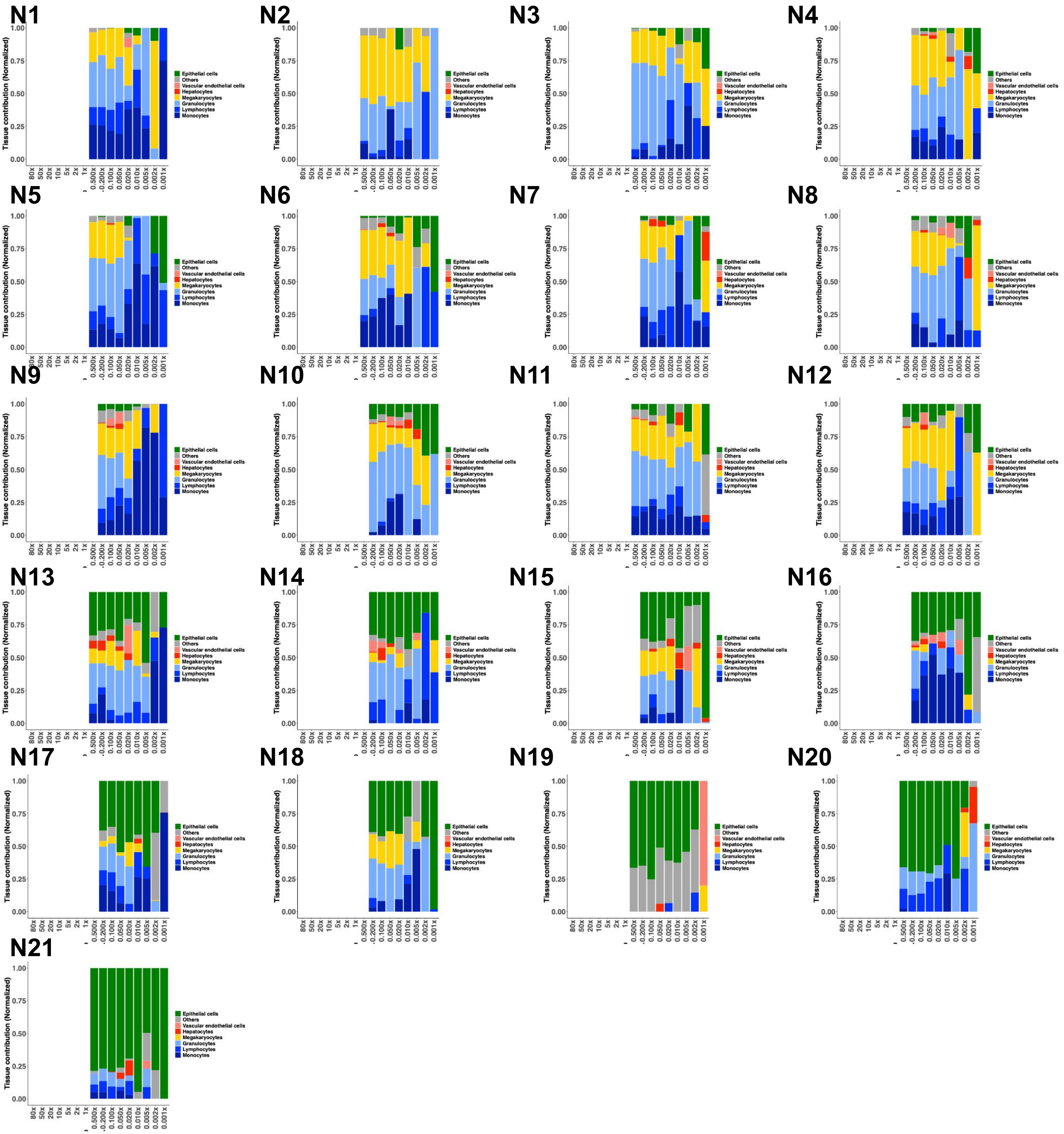

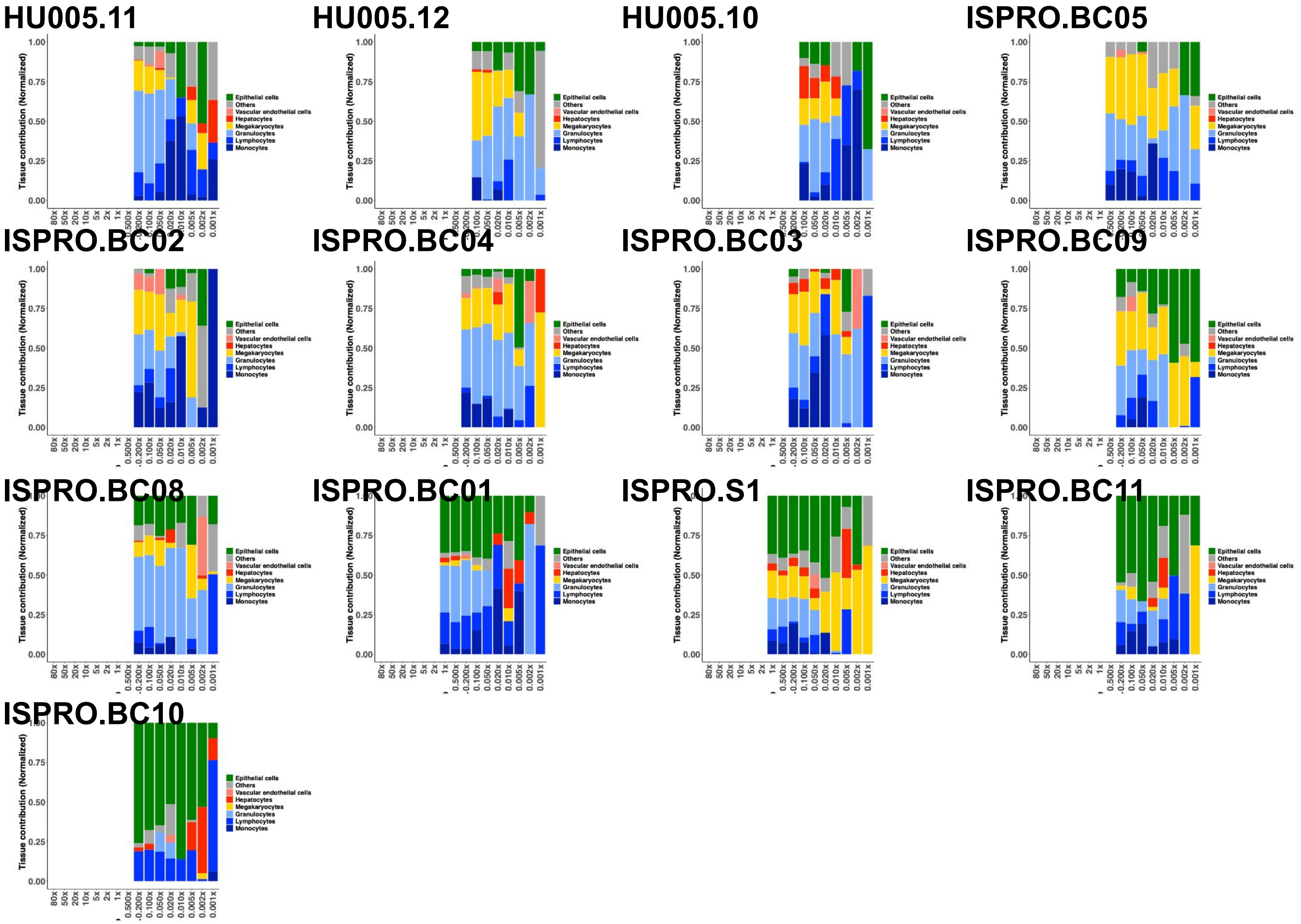

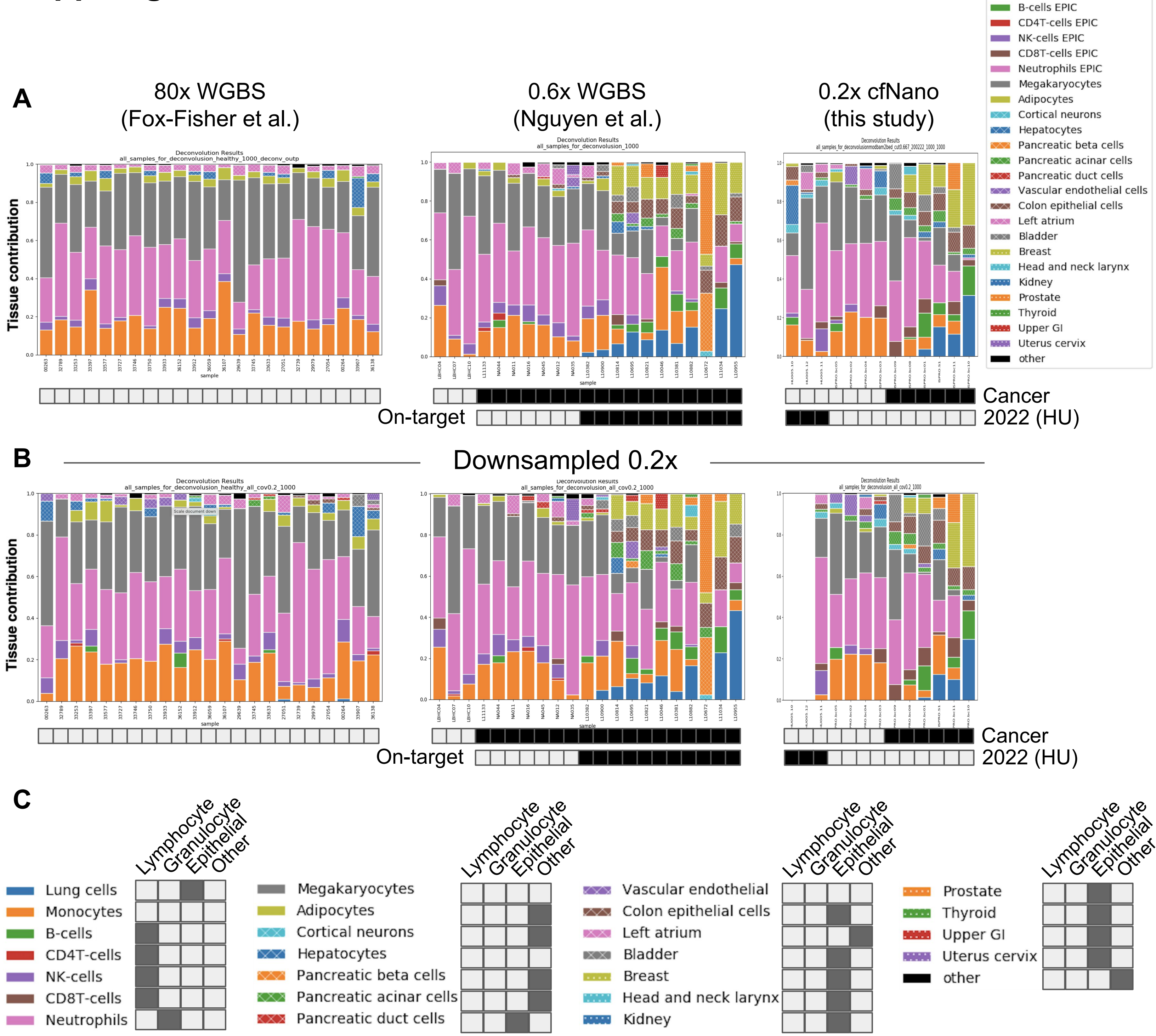

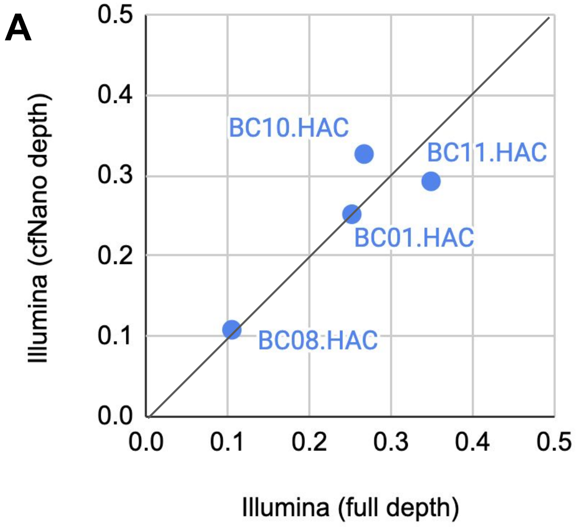

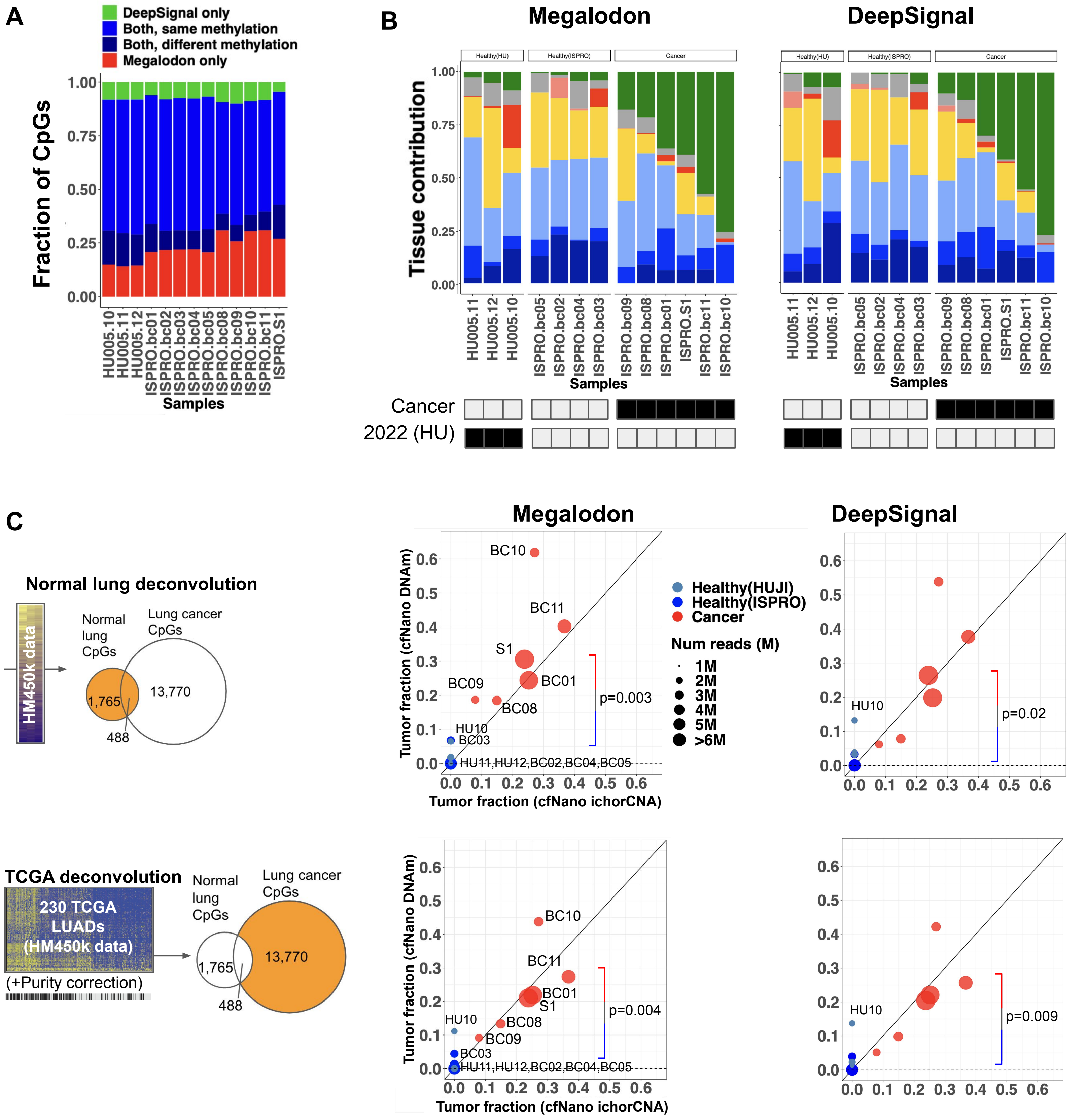

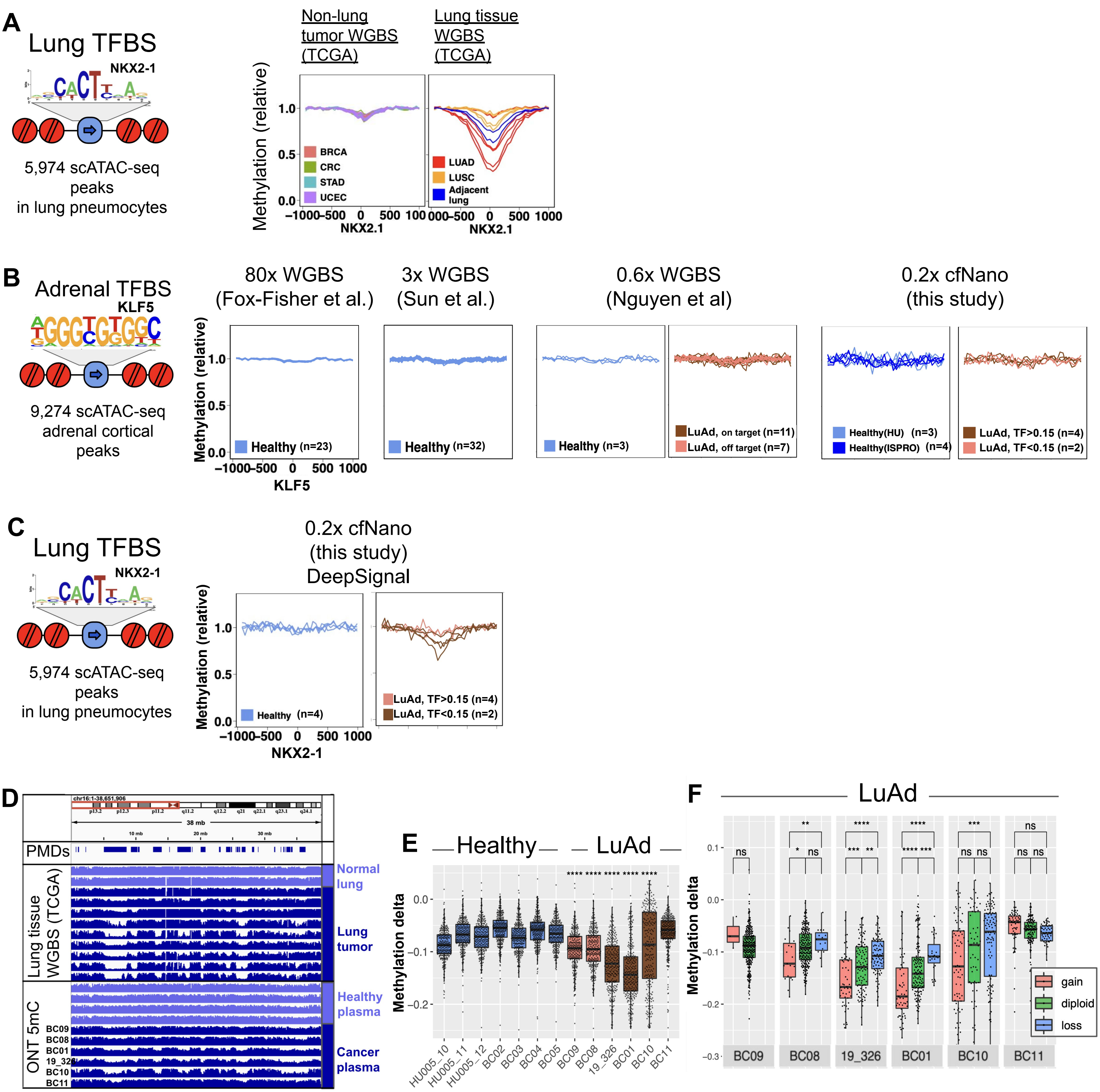

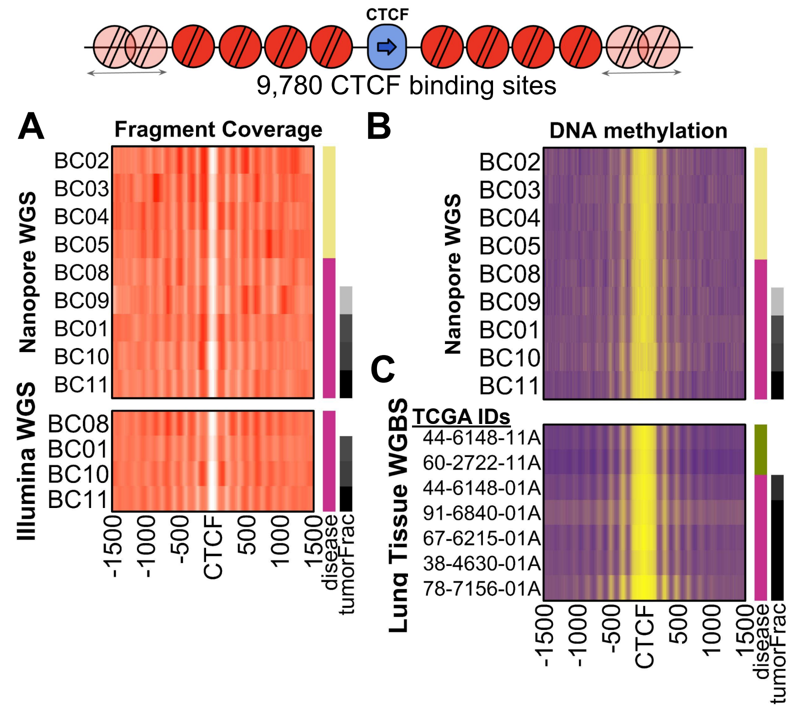

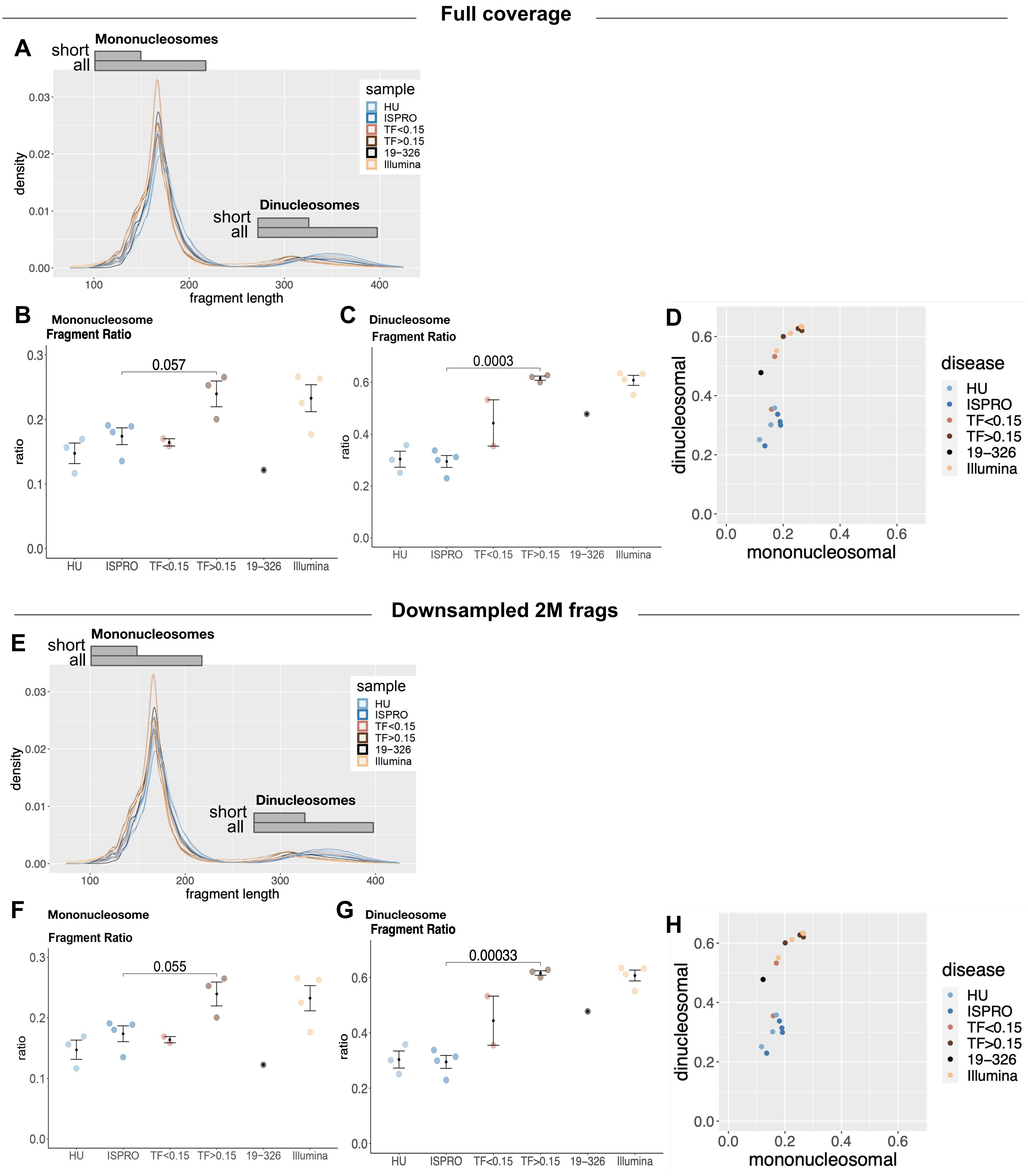

#### Supp. Fig 10

##### Full coverage

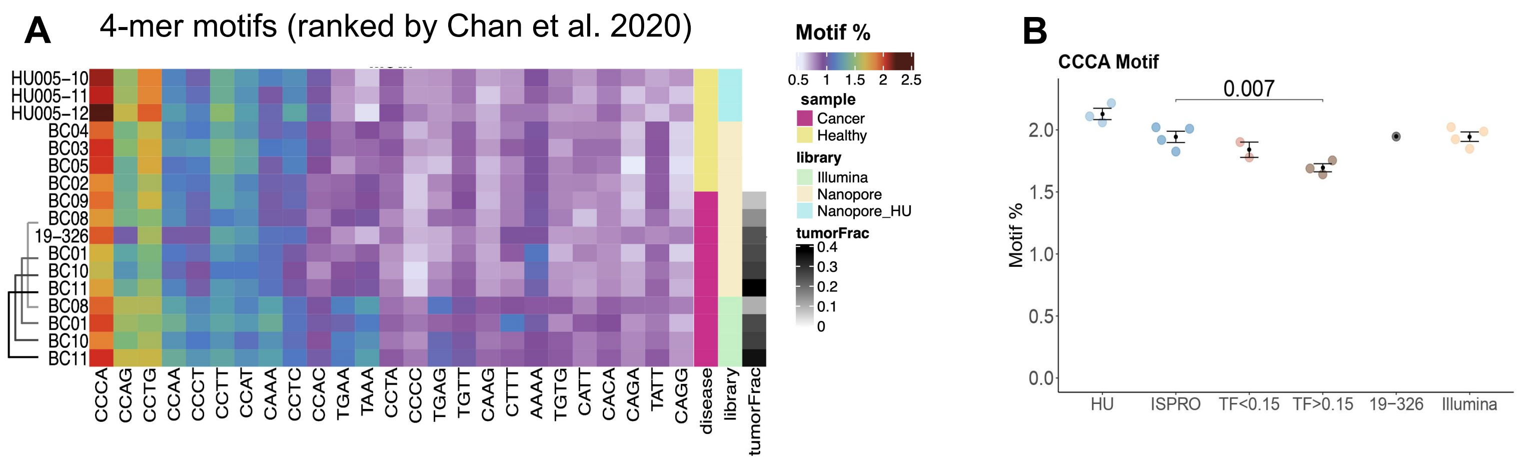


##### Downsampled 2M frags

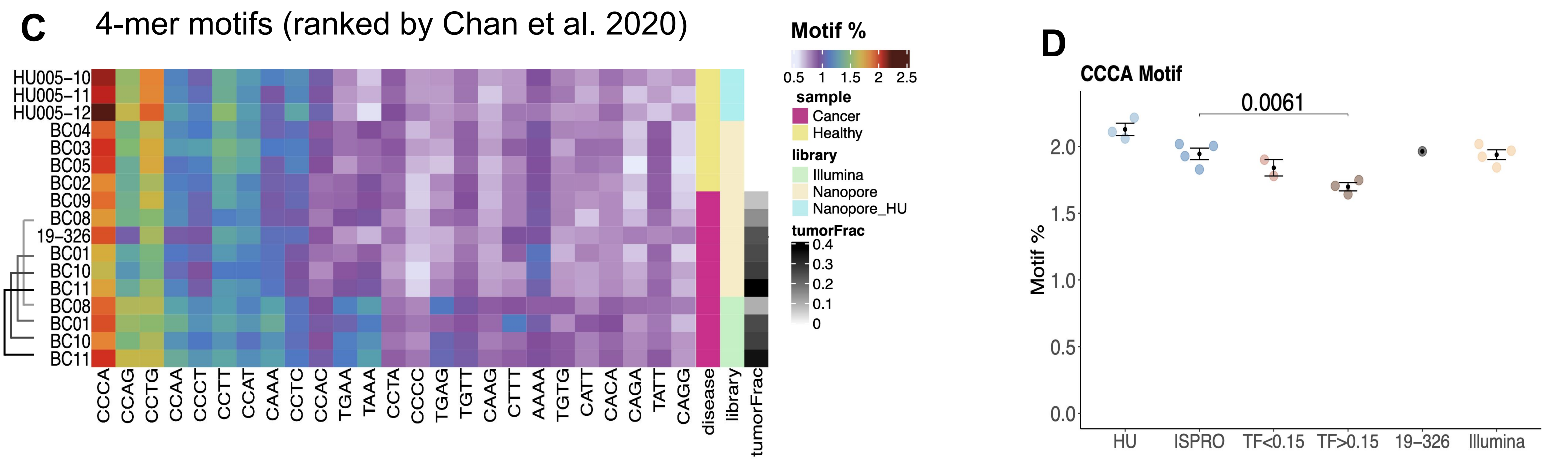
