## Supplementary Data File 1 for "Detecting cell-of-origin and cancer-specific methylation features of cell-free DNA from Nanopore sequencing": ichorCNA-cfNano-original.pdf

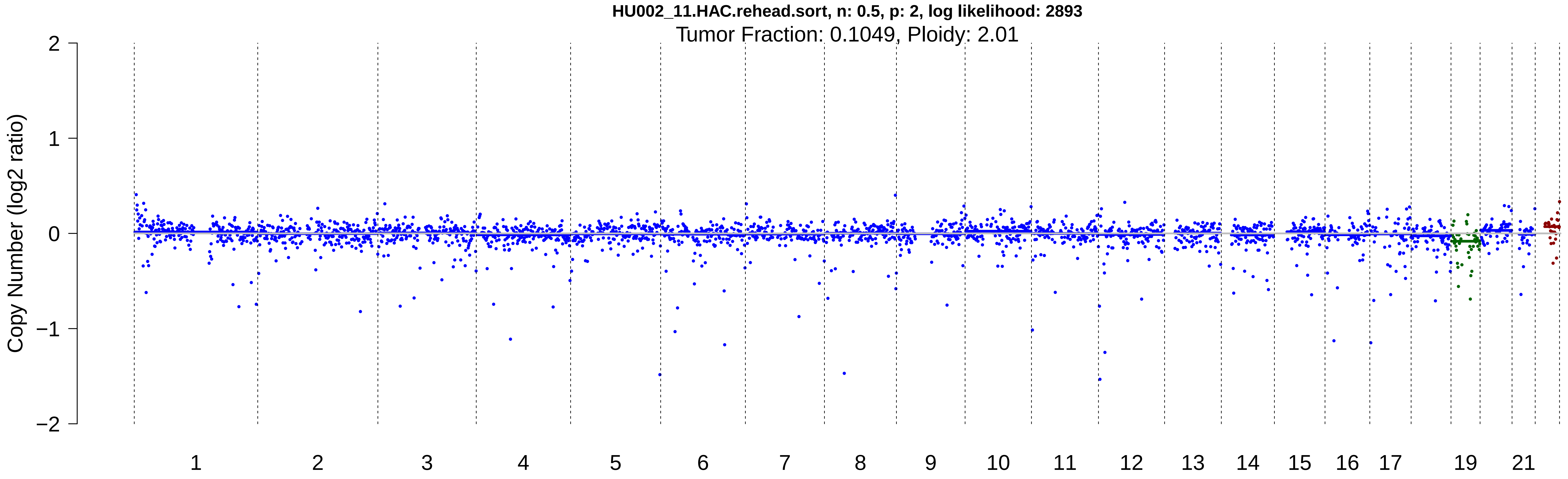

HU002_11.HAC.rehead.sort, n: 0.5, p: 2, log likelihood: 2893
Tumor Fraction: 0.1049, Ploidy: 2.01
Copy Number (log2 ratio)
2
1
0
−1
−2
19
21

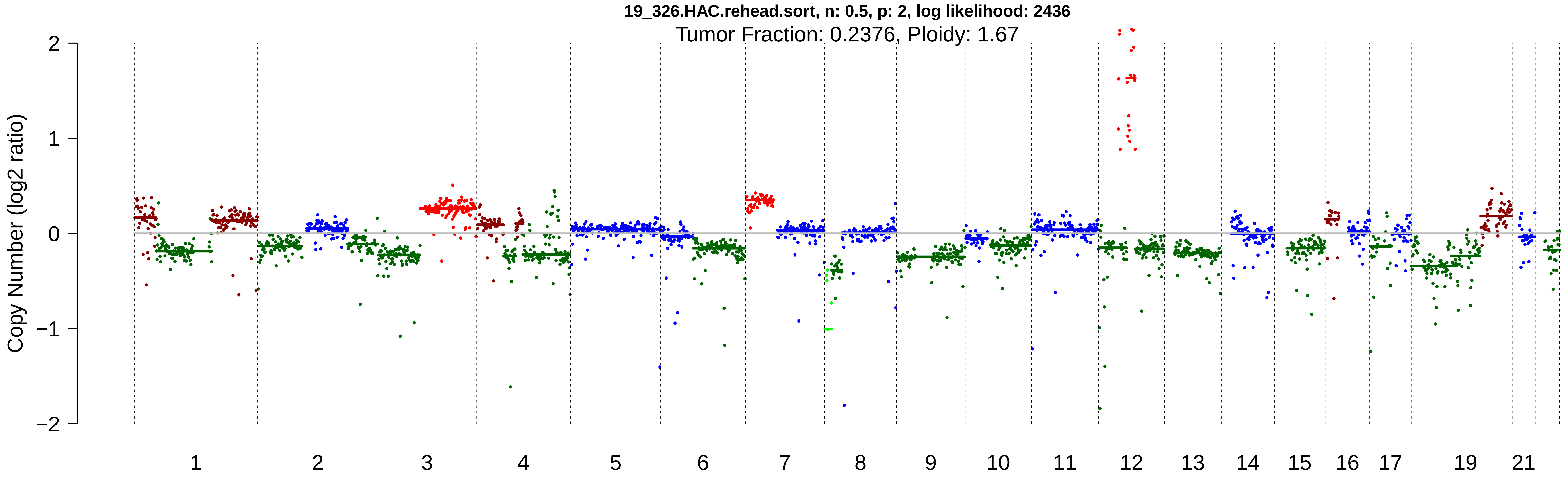

19_326.HAC.rehead.sort, n: 0.5, p: 2, log likelihood: 2436
Tumor Fraction: 0.2376, Ploidy: 1.67
Copy Number (log2 ratio)
2
1
0
−1
−2
19
21

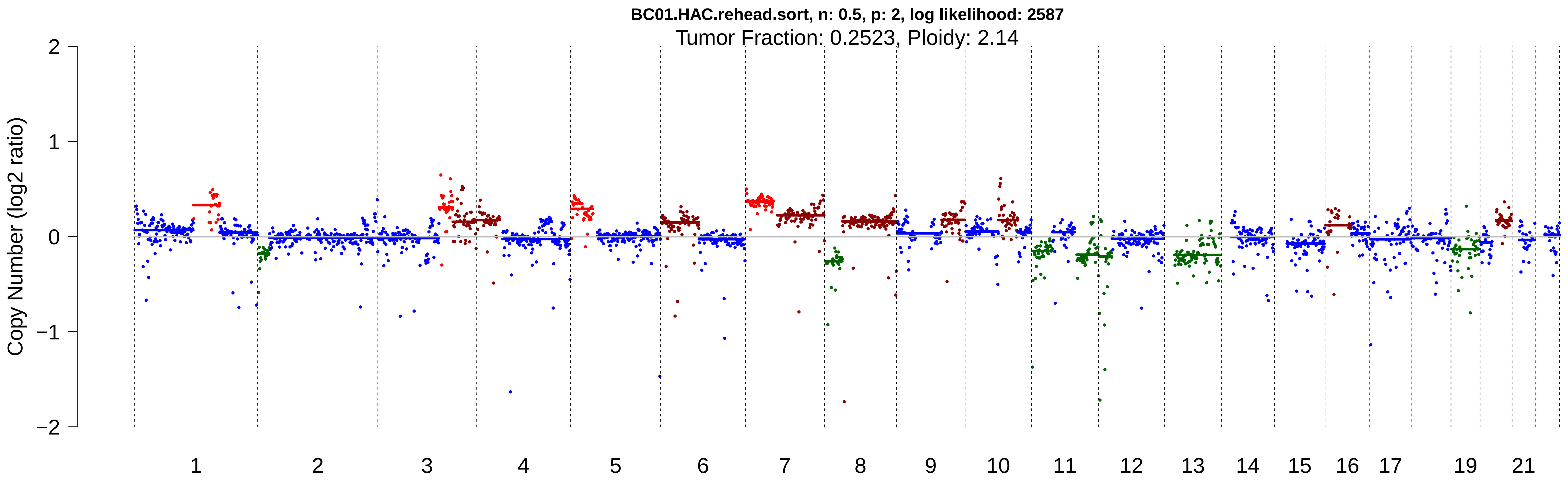

BC01.HAC.rehead.sort, n: 0.5, p: 2, log likelihood: 2587
Tumor Fraction: 0.2523, Ploidy: 2.14
Copy Number (log2 ratio)
2
1
0
−1
−2
19
21

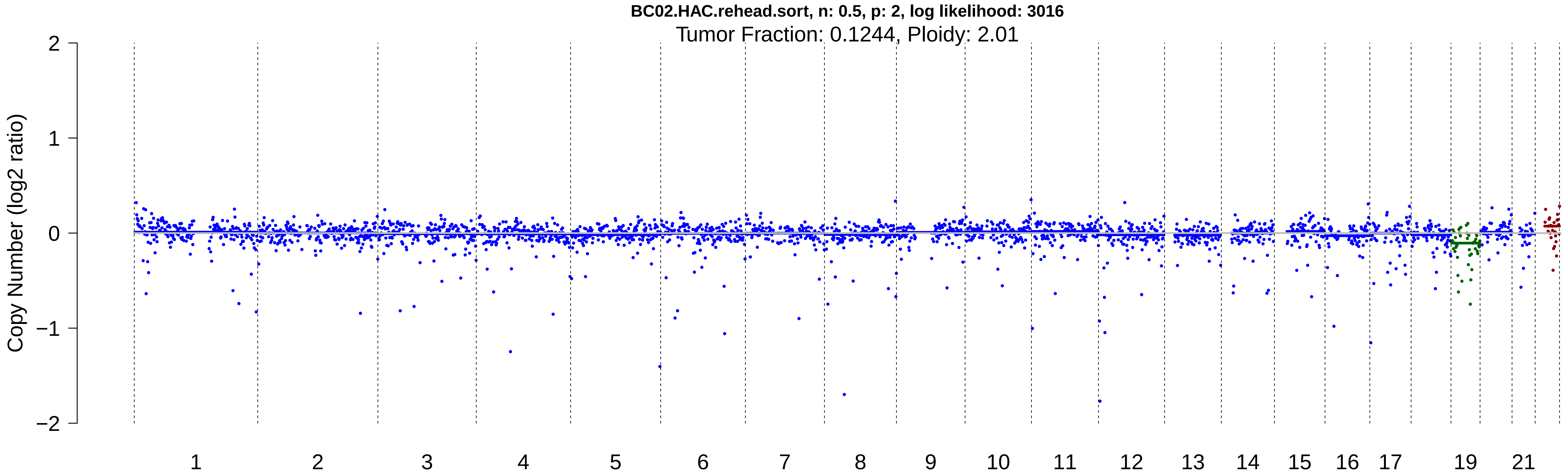

BC02.HAC.rehead.sort, n: 0.5, p: 2, log likelihood: 3016
Tumor Fraction: 0.1244, Ploidy: 2.01
Copy Number (log2 ratio)
2
1
0
−1
−2
19
21

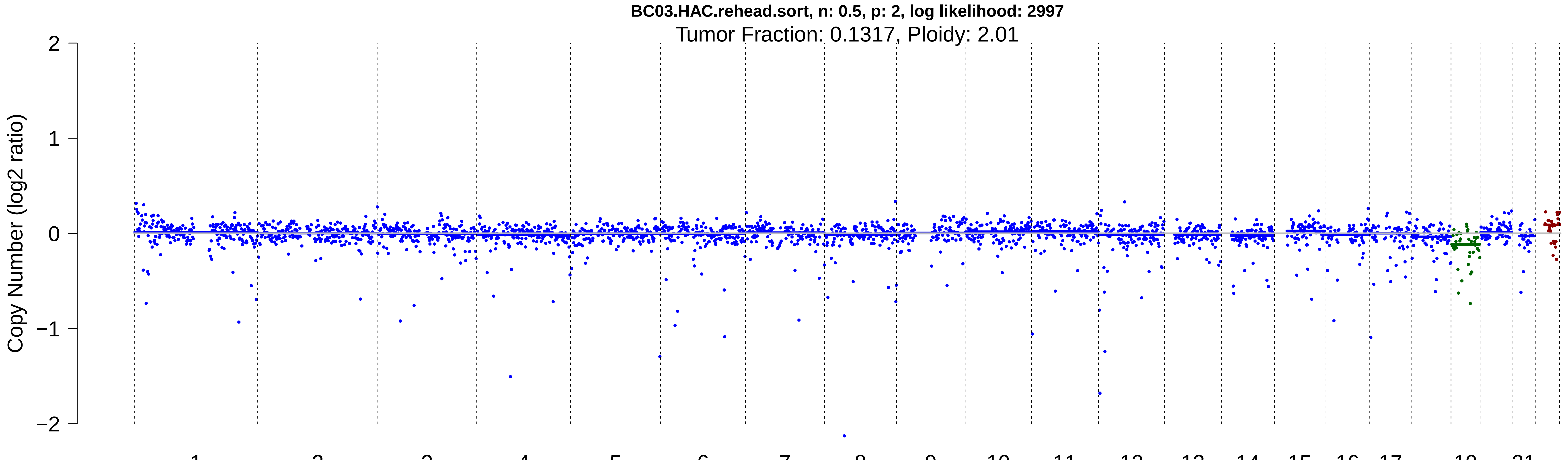
BC03.HAC.rehead.sort, n: 0.5, p: 2, log likelihood: 2997
Tumor Fraction: 0.1317, Ploidy: 2.01
Copy Number (log2 ratio)
2
1
0
−1
−2
19
21

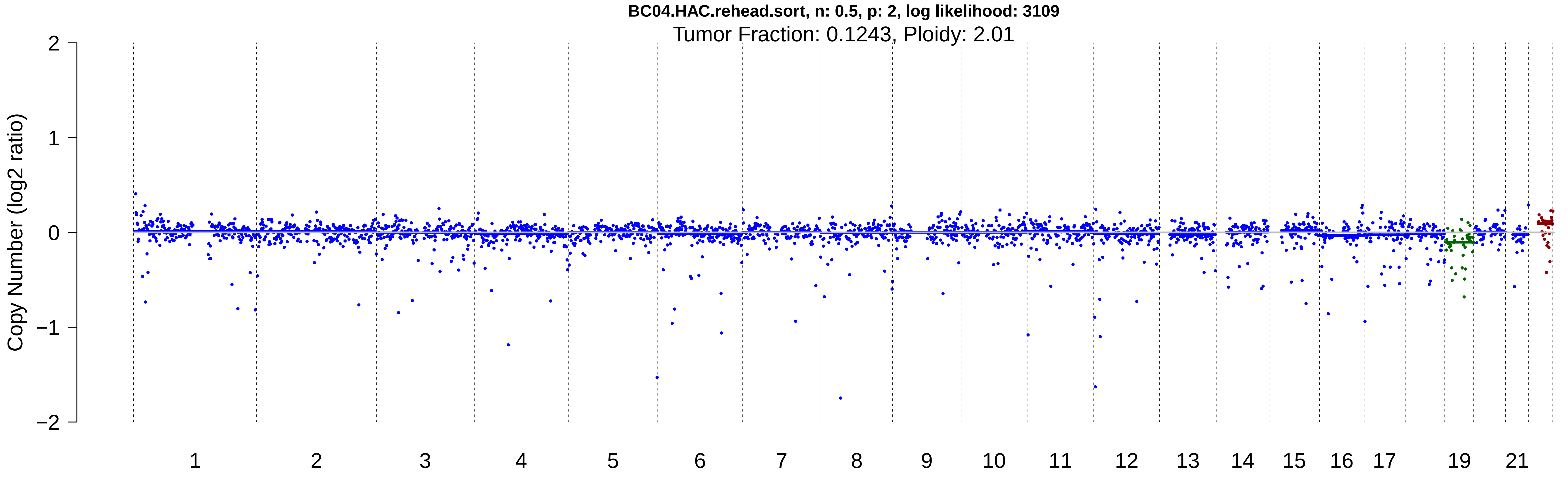

BC04.HAC.rehead.sort, n: 0.5, p: 2, log likelihood: 3109
Tumor Fraction: 0.1243, Ploidy: 2.01
Copy Number (log2 ratio)
2
1
0
−1
−2
19
21

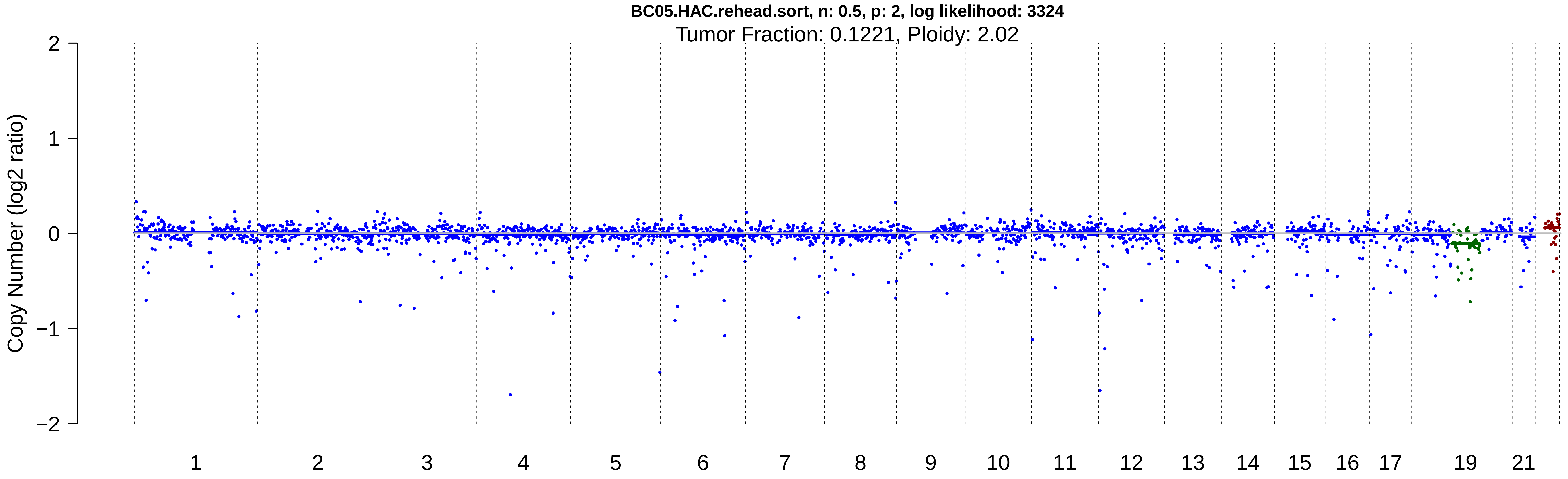
BC05.HAC.rehead.sort, n: 0.5, p: 2, log likelihood: 3324
Tumor Fraction: 0.1221, Ploidy: 2.02
Copy Number (log2 ratio)
2
1
0
−1
−2
19
21

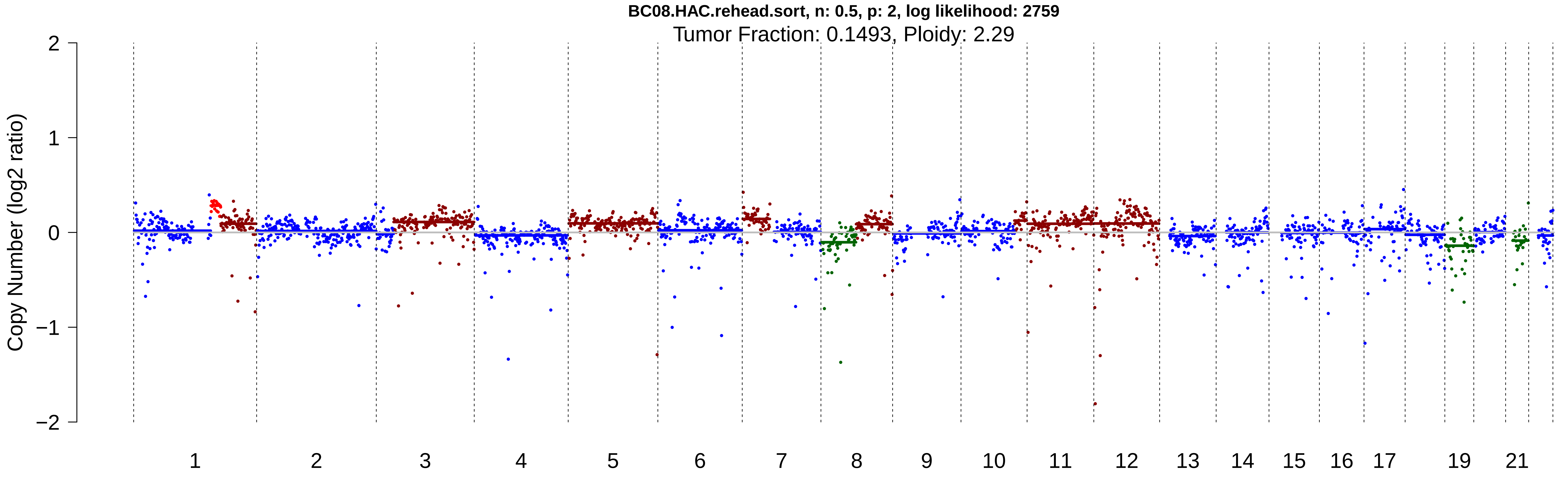

BC08.HAC.rehead.sort, n: 0.5, p: 2, log likelihood: 2759
Tumor Fraction: 0.1493, Ploidy: 2.29
Copy Number (log2 ratio)
2
1
0
−1
−2
19
21

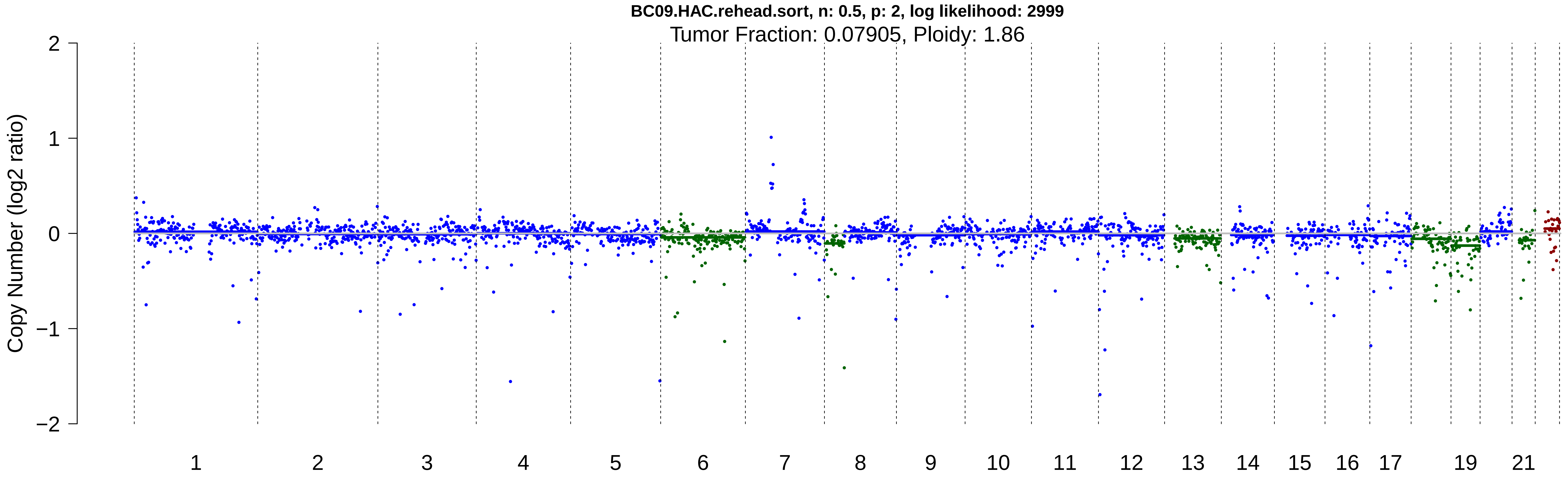

BC09.HAC.rehead.sort, n: 0.5, p: 2, log likelihood: 2999
Tumor Fraction: 0.07905, Ploidy: 1.86
Copy Number (log2 ratio)
2
1
0
−1
−2
19
21

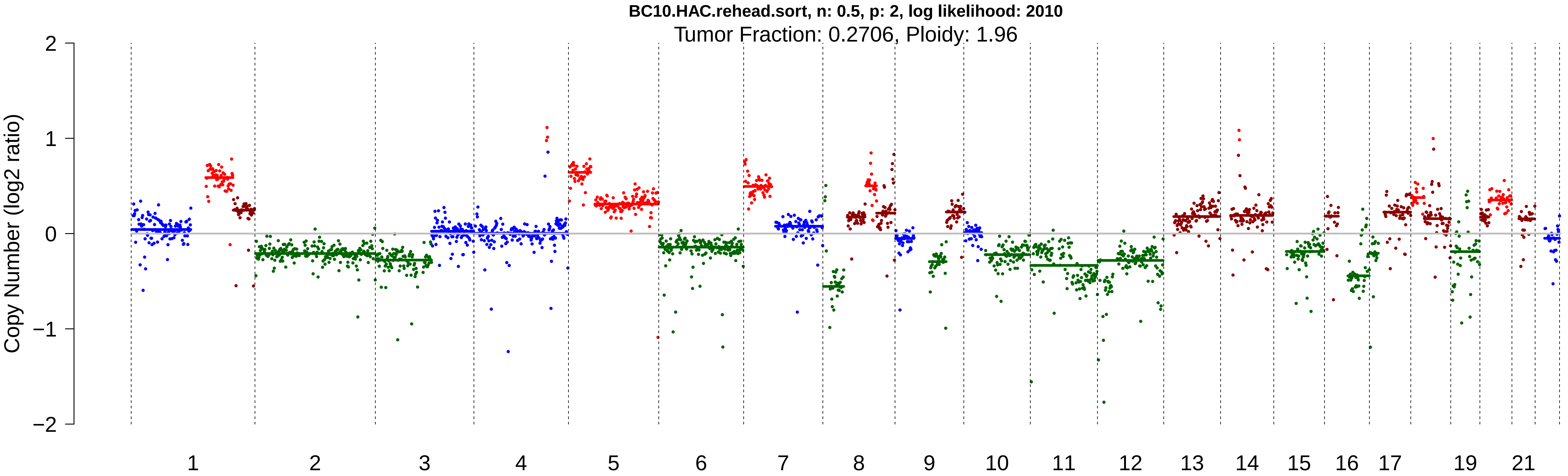

BC10.HAC.rehead.sort, n: 0.5, p: 2, log likelihood: 2010
Tumor Fraction: 0.2706, Ploidy: 1.96
Copy Number (log2 ratio)
2
1
0
−1
−2
19
21

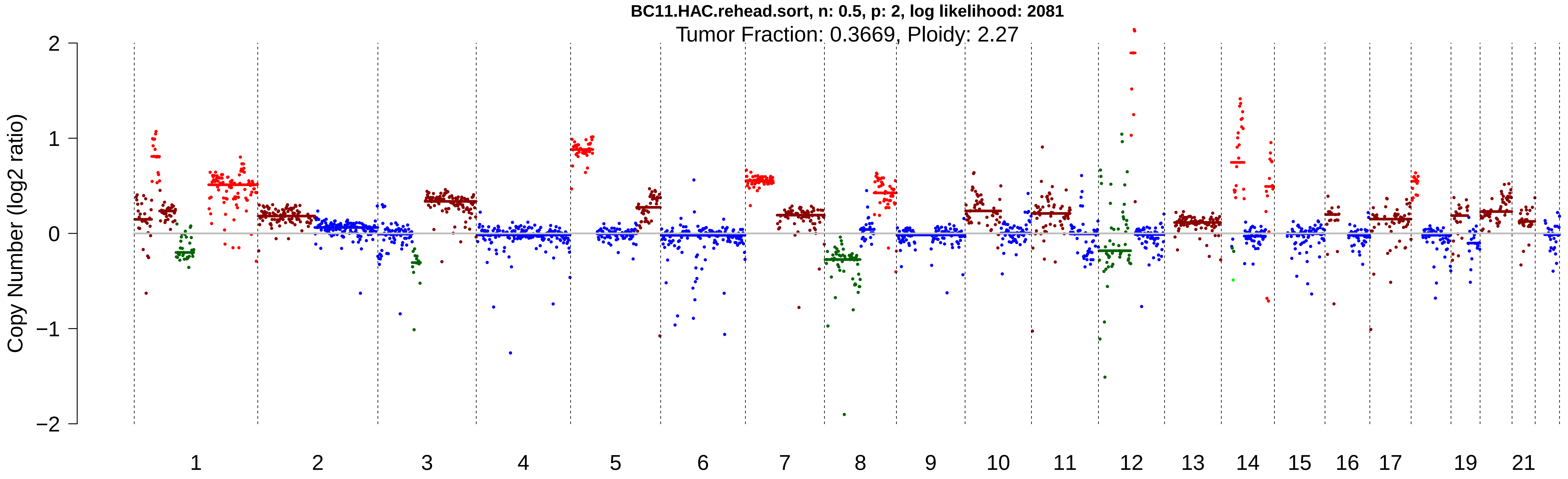

BC11.HAC.rehead.sort, n: 0.5, p: 2, log likelihood: 2081
Tumor Fraction: 0.3669, Ploidy: 2.27
Copy Number (log2 ratio)
2
1
0
−1
−2
19
21

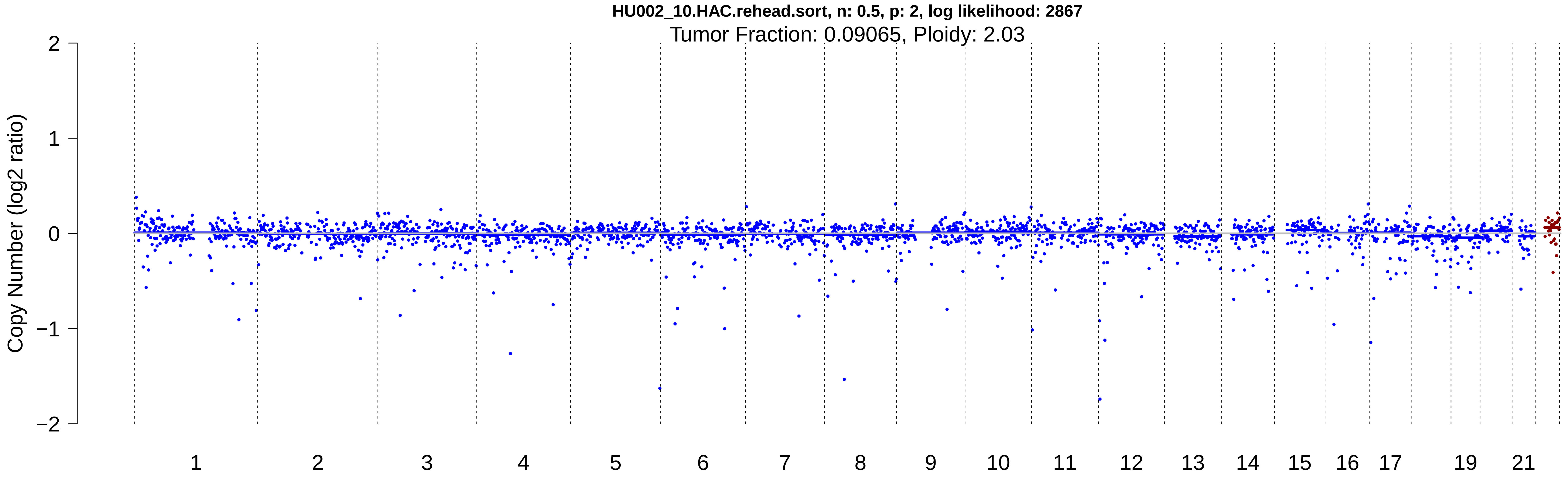
HU002_10.HAC.rehead.sort, n: 0.5, p: 2, log likelihood: 2867
Tumor Fraction: 0.09065, Ploidy: 2.03
Copy Number (log2 ratio)
2
1
0
−1
−2
19
21

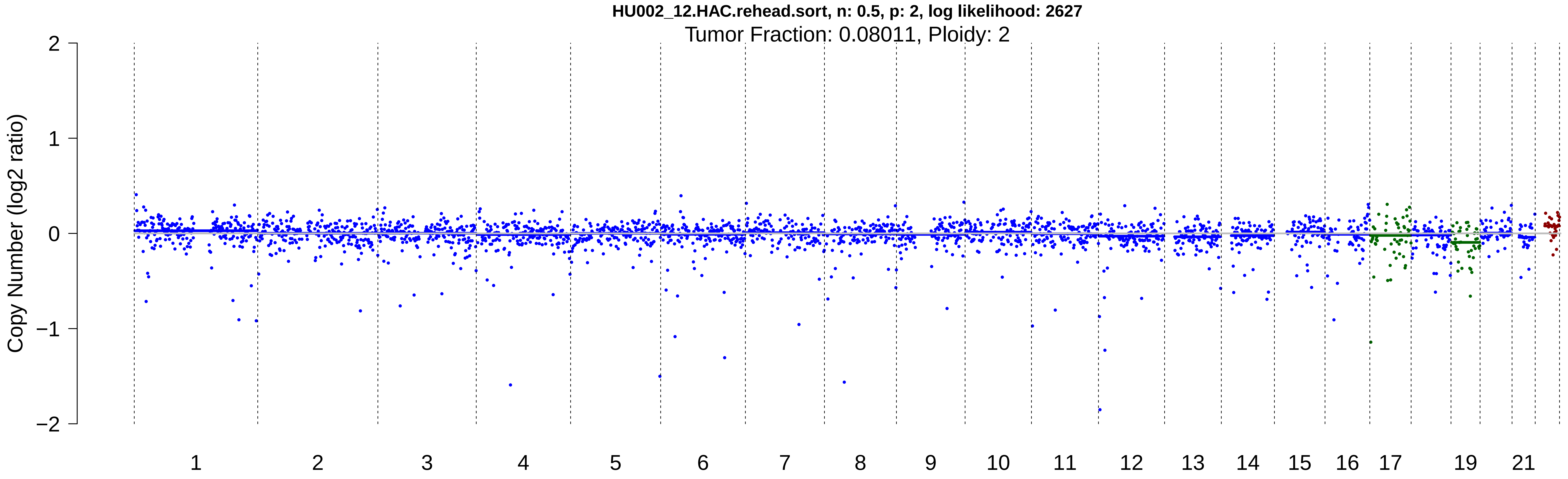
HU002_12.HAC.rehead.sort, n: 0.5, p: 2, log likelihood: 2627
Tumor Fraction: 0.08011, Ploidy: 2
Copy Number (log2 ratio)
2
1
0
−1
−2
19
21
